# Sex chromosomes and chromosomal rearrangements are key to behavioural sexual isolation in *Jaera albifrons* marine isopods

**DOI:** 10.1101/2025.01.08.631900

**Authors:** A Ribardière, C Daguin-Thiébaut, J Coudret, G Le Corguillé, K Avia, C Houbin, S Loisel, P-A Gagnaire, T. Broquet

## Abstract

The lack of sexual attraction between individuals from different populations is a direct barrier to gene flow between these populations. Here we focus on the evolution of this class of isolating mechanism, behavioural sexual isolation, through the empirical study of two closely related species of small marine isopods. The males of *Jaera albifrons* and *J. praehirsuta* similarly engage females in tactile courtship by brushing a specific region of the female’s back, but they do so with divergent sets of specialised setae and spines, and female choice results in strong reproductive isolation. Using bi-allelic SNP genotypes obtained from double-digest RAD sequencing of individuals from natural populations and controlled crosses, we found that secondary contacts between *J. albifrons* and *J. praehirsuta* resulted in different levels of heterospecific gene flow depending on the ecological context. Comparison of the genomic landscapes of differentiation in the two most contrasting situations (extremely low heterospecific gene flow in one region of western France, but strong introgressive hybridisation in another), combined with linkage map analyses, allowed us to conclude that genomic regions resistant to interspecies gene flow are primarily located either on the sex chromosomes or on rearranged chromosomes (several fusion-scissions and one reciprocal translocation). These genomic regions show low recombination, and in two cases QTL analyses found genetic variation associated with several male courtship traits. These results suggest that a long period of allopatry may have allowed the divergent co-evolution of male traits and female preferences, with genetic bases located at least in part in non-recombining regions on sex chromosomes and rearranged chromosomes.

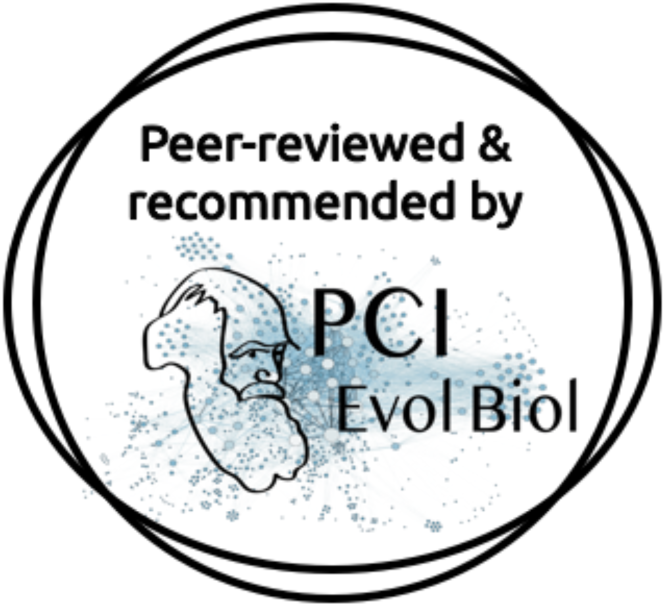

## Introduction

Sexual reproduction, shared by almost all eukaryotes, requires successful gamete transfer between partners and successful syngamy. Any evolutionary divergence between groups of individuals in the way these tasks are accomplished can immediately create reproductive barriers. The resulting effect of such barriers, sexual isolation, seems to have played a major role in speciation, particularly in animals and flowering plants (Mayr 1963; Coyne and Orr 2004; Ellis and Oakley 2016; Janicke et al. 2018; Cally et al. 2021; Shaw et al. 2024). In a recent review of the literature, Shaw et al. (2024) note the rapidity with which sexual isolation can evolve, the strength of this barrier relative to others, and its role in maintaining reproductive isolation between species. While these general conclusions do not underestimate the heterogeneity of situations and the need for further investigation across a wider part of the tree of life, it does emphasise the global importance of sexual isolation in speciation and highlights the need to understand its evolution (Shaw et al. 2024).

One step in understanding the evolution of sexual isolation is to better characterise the genetic basis of sexual barriers and the role of genomic architectures (e.g. Merrill et al. 2024). While sexual phenotypes attributable to multiple interacting loci, each with small, varying allelic effects remain difficult to identify, there are sexual phenotypes involved in reproductive isolation that are mediated by a few genes or genomic regions of large effect (e.g. for courtship behavior, Arbuthnott 2009). Technological advances are making it increasingly possible to characterise this genetic determinism in detail (Fan et al. 2013; e.g. Poelstra et al. 2014; Ding et al. 2016; Enge et al. 2021). A less direct, more general approach, describing the genomic distribution of genetic differentiation between species, has opened a potential window on the semi-permeability of genomes to interspecific gene flow (reviewed in Seehausen et al. 2014; Ravinet et al. 2017). In particular, such studies have shown that highly differentiated genomic regions are often associated with low recombination, an observation compatible with the role of recombination arrest in preventing mixing of genetic material at barrier loci and maintaining allelic associations between loci (Felsenstein 1981).

Chromosomal rearrangements, which are frequently observed between closely related species (White 1973; Searle 1993; Augustijnen et al. 2024), have been singled out as a potentially highly effective cause of recombination suppression in the context of speciation, especially with respect to the evolution of sexual isolation (Trickett and Butlin 1994). Chromosomal inversions, in particular, have been shown in many genomic studies to play a central role in reproductive isolation (Noor et al. 2001b; Rieseberg 2001; Faria and Navarro 2010; Zhang et al. 2021; Berdan et al. 2024; Le Moan et al. 2024). Similarly, sex chromosomes represent another region of the genome associated with potentially drastic modifications of the recombination landscape that can have profound consequences on the evolution of reproductive isolation (Qvarnstrom and Bailey 2009; Beukeboom and Perrin 2014; Payseur et al. 2018; Fraïsse and Sachdeva 2021).

In accordance with these hypotheses, recent empirical studies have yielded a substantial body of results indicating that differentiation on the sex and/or rearranged chromosomes is typically greater than that observed on the rest of the genome in a multitude of taxa (Presgraves 2018). However, there are several important issues to consider when interpreting these patterns. A first serious challenge is to disentangle the direct effects of sex chromosomes and chromosomal rearrangements on the fitness of heterokaryotic hybrids from their indirect effects through recombination arrest (Searle 1993; Trickett and Butlin 1994; Noor et al. 2001b; Rieseberg 2001; Faria and Navarro 2010; Lucek et al. 2023; Berdan et al. 2024). Hybrids that are heterozygous for structural variants such as large inversions, fusions and translocations can suffer reduced fitness if pairing, chiasma formation or segregation does not proceed properly at meiosis, resulting in germ cell death or unbalanced (aneuploid) gametes (White 1978; Searle 1993; Faria and Navarro 2010). The occurrence and extent of these processes can be difficult to assess.

Another well-known problem is that the interpretation of genomic differentiation patterns in terms of interspecific gene flow is complicated by confounding factors such as local variation in effective size, mutation rate, and local gene density, some of which are particularly acute in regions of low recombination (Burian et al. 1988; Noor and Bennett 2009; Cruickshank and Hahn 2014; Seehausen et al. 2014; Ravinet et al. 2017). One way out of this conundrum is to identify the effect of gene flow on the genomic landscape of differentiation between pairs of populations that are thought to differ in the amount of gene flow that occurs (typically pairs of populations in sympatry vs. allopatry, Noor and Bennett 2009; Sætre and Ravinet 2019), and to overlay this with information on the distribution of recombination.

The present study focuses on a pair of closely related intertidal marine isopods (*Jaera albifrons* Leach, 1814 and *J. praehirsuta* Forsman, 1949, Fig. 1) for which i) sexual isolation based on courtship behaviour is strong and has been a critical component of speciation (Solignac 1981; Ribardière et al. 2021), ii) Robertsonian chromosomal polymorphism is extensive both within and between species and does not seem to generate post-zygotic isolation (Staiger and Bocquet 1956; Lécher 1967b; Lécher and Prunus 1971; Solignac 1978; Ribardière et al. 2021), iii) sex is determined by a female-heterogametic genetic system with a large metacentric Z chromosome and two smaller acrocentric W1 and W2 chromosomes (Staiger and Bocquet 1954; Lécher 1967b) and iv) low coverage genetics suggested sharp contrasts in the permeability to gene flow along their approximately 0.9 Gb genome (Mifsud 2011; Ribardière et al. 2017).

**Figure 1 –.**
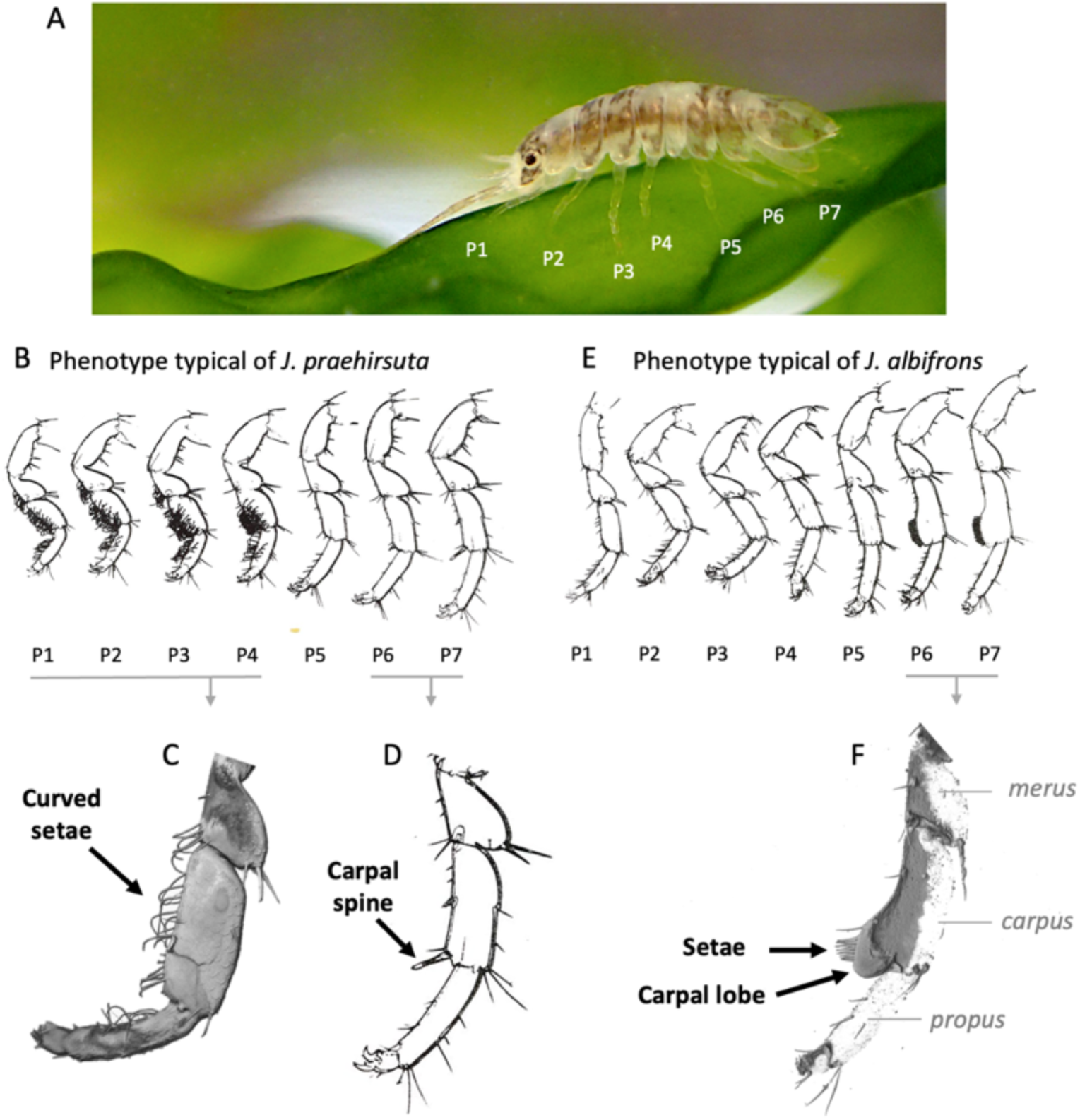
Sexual phenotypes of *Jaera praehirsuta* and *J. albifrons* males. A) An adult individual showing the seven pairs of legs which, in males, bear the secondary sexual characteristics used for courtship (photograph credit: G. Evanno & T. Broquet). The sexual phenotype typical of *J. praehirsuta* males (B), is composed by curved setae (C) on the *propus, carpus*, and *merus* segments of the first four peraeopods ("legs" P1-P4), and one or two rigid spines (D) on the *carpus* of the posterior peraeopods (P6-7). In contrast, male *J.albifrons* (E) use brushes formed on P6 and P7 by straight setae set on a carpal lobe (F) and have no sexual setae on P1-4. A range of intermediate phenotypes may form in case of hybridization. The drawings in B, D, and E are reproduced from Solignac (1981) with authorization. The close-up pictures in C and F were obtained with a confocal laser scanning microscope and processed using software Fiji and Imaris (photograph credit to Sébastien Colin).

Most importantly, these two species co-occur in sympatry in different geographic regions where ecological variation leads to either strong isolation or introgressive hybridization, but with consistently high behavioural sexual isolation (Solignac 1969; Ribardière et al. 2021). In Brittany, western France, *J. albifrons* is mainly found under pebbles and stones while *J. praehirsuta* is mainly found on brown algae, amounting to 98% ecological isolation even in areas where the two micro-habitats are directly in contact (Ribardière et al. 2021). By contrast, further north and east along the coast (region Normandy), where there are less sheltered fucoid belts and algae are not used as habitat, the two species coexist under pebbles and stones (no ecological isolation). However, lab experiments have shown that sexual isolation, driven by male courtship and female choice, is intrinsically strong in both regions. In nature, this barrier can operate effectively only if there is no ecological barrier forbidding heterospecific encounters, as in Normandy, where lab experiments estimated sexual isolation to 92% (Ribardière et al. 2021).

We can thus contrast two situations involving locally mixed populations: one (Brittany) where the two species are fully or nearly fully reproductively isolated due to the combination of ecological and sexual barriers, and another (Normandy) where the two species are strongly but incompletely isolated by sexual isolation only, allowing for ongoing introgressive hybridization (Solignac 1969; Ribardière et al. 2017, 2021). This system is interesting because we can compare the genomic landscape of differentiation under contrasting levels of interspecies gene flow. This provides an opportunity to investigate the impact of gene flow on the genomic landscape of differentiation between species that are likely to differ in their chromosome structure and genomic landscape of recombination.

Here we used restriction-associated DNA sequencing (RADseq) of individuals from natural populations and controlled crosses to investigate the signature of sexual isolation on genetic differentiation between *J. albifrons* and *J. praehirsuta*. Our specific aims were as follows.

1. To estimate genome-wide genetic differentiation between species in several sympatric population pairs on the west coast of France, with the aim of refining the spatial variation in interspecies gene flow documented by Solignac (1969) and Ribardière et al. (2017), and inferring historical scenarios of population divergence that may have produced these contrasting situations.
2. To document broad patterns of the genomic recombination landscape within each species and chromosomal polymorphism between species by constructing several linkage maps.
3. To investigate, using a quantitative trait locus (QTL) approach, how the genetic determinism of male sexual traits involved in courtship is linked to the major features of genome architecture identified in point 2.
4. Finally, to identify genomic regions associated with behavioural sexual isolation by examining how interspecies gene flow shapes the genomic landscape of differentiation, building upon the spatial variation in gene flow documented in (1), the recombination landscape and chromosomal polymorphisms described in (2), and the genomic regions associated with male sexual traits identified in (3).

## Material and methods

### Species

*J. albifrons* and *J. praehirsuta* both belong to the *Jaera albifrons* complex, a small group of five small (2-5 mm) isopod species that live on the shores of the cold and temperate coasts of the North Atlantic Ocean and adjacent seas, ranging from southern Greenland in the north to Portugal and the north-eastern United States in the south (Bocquet 1953; Solignac 1978). These species have largely overlapping distributions in the high intertidal zone, where they may occupy different ecological niches that vary in intertidal position, exposure, salinity and substrate (Naylor and Haahtela 1966; Jones 1972). Yet these ecological differences are not stable or strict, and sexual isolation through mate choice mechanisms is decisive for reproductive isolation between species (Solignac 1981; Ribardière et al. 2021). Sexual isolation is achieved through a tactile courtship display in which males use sexual setae and spines to brush a specific area of the female’s back, which can then accept or reject intercourse. While this general courtship behaviour is the same across species, males differ in the specific distribution of sexual setae and spines, thus proposing divergent tactile stimuli in conjunction with divergent female preferences (Bocquet 1953; Solignac 1981).

### Sampling

Individuals were sampled at 11 sites along the Channel coast between 2014 and 2016 (Fig. 2A and supplementary material Table S1) as described in Ribardière et al. (2017). Each individual was sexed and males were identified under a dissecting microscope within a few hours after collection and then conserved in ethanol. Individuals with secondary sexual characteristics typical of both *J. albifrons* and *J. praehirsuta* were classified as "intermediate phenotypes", while all other individuals were assigned to one species. Intermediates (“hybrid types” 5-13 in Solignac 1978; and Fig. 1 in Ribardière et al. 2017) typically have both a few curved setae on the anterior pereiopods (similar to Fig. 1C) and a visible carpal lobe bearing an altered brush or spine on the posterior pereiopods (intermediate between Fig. 1D and F). Only adult males were used in the analyses presented here (except for linkage maps, as described below). In addition, nine *J. ischiosetosa* Forsman, 1949 males used as an out-group came from randomly selected locations along the coast between sites 1 and 6 (see supplementary material Fig. S1).

**Figure 2 –.**
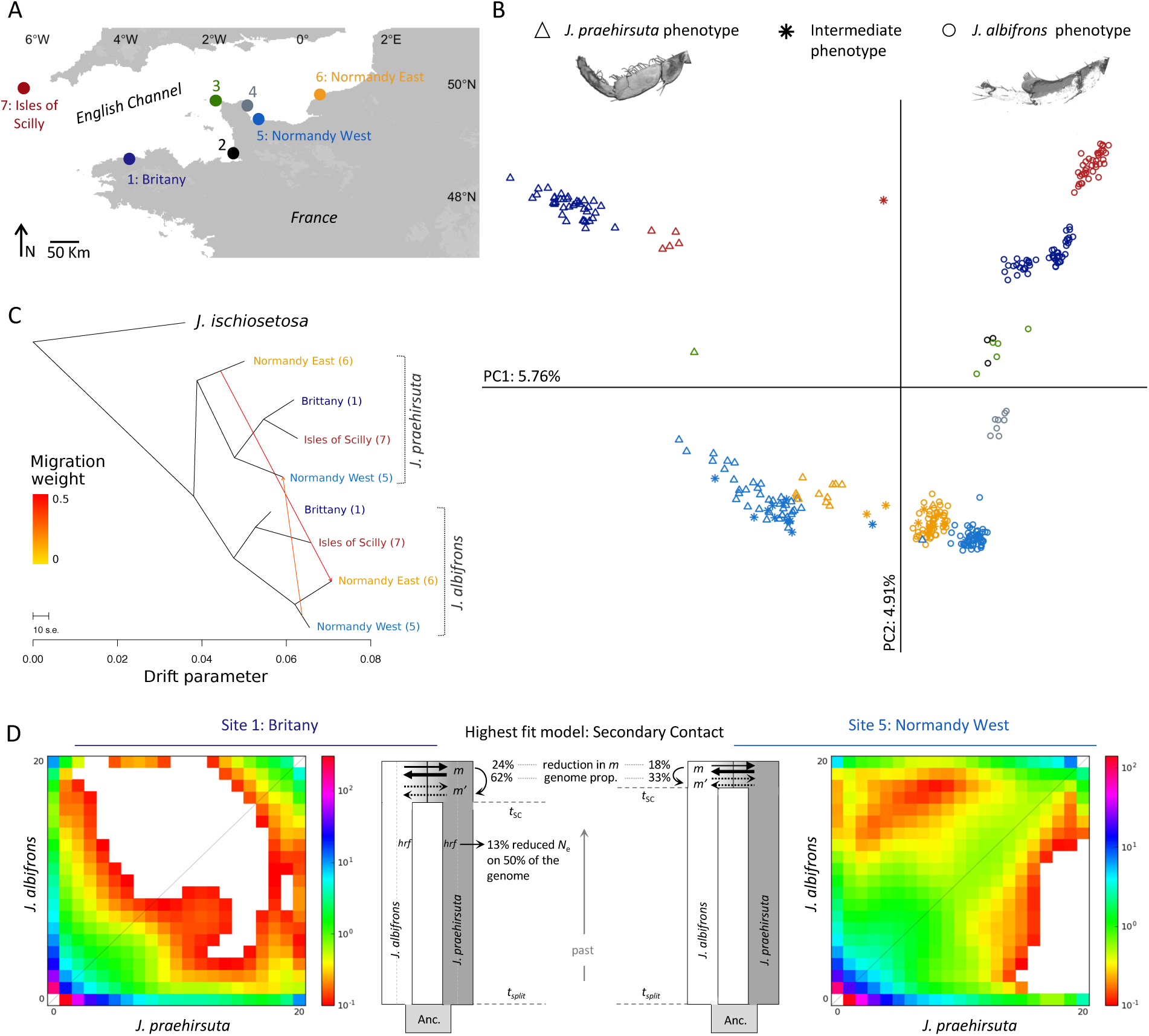
A) Sampling sites *for J. albifrons* and *J. praehirsuta.* Some analyses were restricted to sites with sufficient individuals of each species: 1 (Britany), 5 (Normandy West), 6 (Normandy East), and 7 (Isles of Scilly). B) Principal component analysis based on 9718 SNPs derived from ddRAD-seq. Axis 1 separates individuals according to species, with individuals bearing intermediate phenotypes clustering either together with individuals of one of the two species or, more rarely, in between the two species. Axis 2 correlates with the geographical origin of the samples (numbers and colours as in map). C) Best-fit tree inferred by TreeMix for four mixed *J. albifrons* / *J. praehirsuta* populations allowing for two migration events (represented by coloured arrows). The first inferred split separates all *J. praehirsuta* from all *J. albifrons* populations, but interspecific gene flow from *J. albifrons* to *J. praehirsuta* is detected by TreeMix in site 5 (Normandy West), and in the opposite direction in site 6 (Normandy East). Allowing for two migration events resulted in a better fit than without migration (see Fig. S3). D) Joint allelic frequency spectra generated by *δ*a*δ*i for 1913 SNPs in region Brittany (site 1) and 1082 SNPs in region Normandy West (site 5). The colour in each square represents the number of SNPs with allelic frequencies in *J. praehirsuta* and *J. albifrons* as indicated by the x and y axes (on a scale of 0 to 20 copies of the ancestral allele). A simplified representation of the demographic model that provided the best fit to observed genetic data in the *δ*a*δ*i analyses is shown next to each frequency spectrum. In both cases, the observed data were best matched by a secondary contact scenario in which *J. albifrons* and *J. praehirsuta* recently resumed gene flow relative to the inferred time of split. The two models assumed two classes of gene flow (the percentages indicate the proportion of the genome where gene flow is restricted, and the intensity of this restriction), and, only in the case of Brittany (left), heterogeneity in effective population size across the genome (*hrf*). The time proportions are respected in the figure, but the other parameters are only schematic (see text).

### Genotyping

All genetic analyses were based on bi-allelic single nucleotide polymorphism (SNP) genotypes obtained from double-digest RAD-sequencing without reference genome (Peterson et al. 2012; Daguin-Thiébaut et al. 2021). Methods are described in the supplementary material. Briefly, the libraries were sequenced as 125 bp single end reads followed by alignment and SNP calling performed in Stacks v2.52 (Catchen et al. 2011, 2013). The data generated by Stacks were further filtered in *R* v3.5.3 (R Core Team 2019) to remove SNP loci and individuals with more than 10% to 50% of missing data depending on the objective (see supplementary material table S2). Finally, the rate of genotyping error was estimated by counting the number of genotypes that differed within each of the 14 replicate pairs that were included in the library construction and sequencing.

### Genetic structure

The distribution of genetic variation across species and populations was analysed using 9718 SNPs genotyped for 221 *J. albifrons*, 103 *J. praehirsuta*, and 25 individuals with *J. albifrons*-*J. praehirsuta* intermediate phenotypes (Table S1). We also included 9 *J. ischiosetosa* males as out-group (based on the phylogeny reported by Mifsud 2011). Following preliminary analyses, locations 1 (Brittany) and 5 (Normandy West) included individuals from 2 to 3 distinct sites (supplementary material Fig. S1) that were grouped together based on spatial proximity, low genetic differentiation within each species, and identical patterns of differentiation between species, following Ribardière et al. (2017, 2021).

The genetic structure was first explored using principal component analyses (PCA) built with the *R* package *adegenet* (Jombart 2008; Jombart and Ahmed 2011). We then used functions from *dartR* (Mijangos et al. 2022) to estimate genetic differentiation between populations (*F*_ST_) and decide how samples should be grouped for downstream analyses. These population-level statistics were estimated for the four locations with enough individuals sampled in both species (Brittany: site 1, Normandy West and East: sites 5 and 6, and Isles of Scilly: site 7, see Table S1 and Fig. 2A). Results from these preliminary analyses suggested that the genetic differentiation between species varied along the study area (see Results). We thus used Pickrell and Pritchard’s TreeMix approach (Pickrell and Pritchard 2012) to infer the patterns of population splits while allowing for mixture between these populations. TreeMix was set up using *J. ischiosetosa* as root, no sample size correction, and five migration conditions (allowing for 0, 1, 2, 3, or 4 migration events between lineages, such as to allow for admixture between species in each region, and between regions). Thirty runs were performed for each condition to assess the robustness of the trees inferred. The relative goodness of fit of each tree was assessed by comparing the distribution of residuals, the log-likelihood, and the fraction of variance explained by each model (following Pickrell and Pritchard 2012).

### Demographic history of differentiation

The distribution of genetic variation and evolutionary relationships between populations as inferred by PCAs, *F*_ST_estimates, and TreeMix suggested that there were heterogeneous patterns of gene flow between species in different locations (Results). The likelihood of different plausible scenarios for the history of differentiation between *J. albifrons* and *J. praehirsuta* was thus evaluated using the modified version of *δaδi* (Gutenkunst et al. 2009) described in Rougeux et al. (2017). This method includes the ability to model semipermeable migration across the genome due to localized barrier effects and variation in effective population size caused by selection at linked sites. The following alternative models were considered (supplementary material Fig. S4): i) strict isolation (SI), where an ancestral population split into two populations that then diverged without gene flow, ii) isolation with migration (IM), where gene flow occurs at a constant rate during population divergence, iii) ancient migration (AM), where gene flow is discontinued after an initial period of divergence with gene flow, and iv) secondary contact (SC), where gene flow resumes after an initial period of divergence without gene flow. The three models with migration (IM, AM, and SC) were tested with one or two classes of loci for migration rates (therefore allowing or not for heterogeneous migration across the genome) and AM and SC were also tested with or without genomic variation in effective population size (that is, 13 models overall, Fig S4).

The method implemented in *δ*a*δ*i uses a diffusion approximation approach to estimate the joint allelic frequency spectrum (JAFS) predicted to result from each demographic scenario, which can then be compared with the observed JAFS for the populations of interest. We built JAFS for *the J. albifrons* / *J. praehirsuta* pair independently in two locations with contrasted patterns of genetic differentiation and where we had enough individuals (Brittany = site 1 in Figure 2A: 50 *J. albifrons* and 39 *J. praehirsuta;* Normandy West = site 5: 40 *J. albifrons* and 38 *J. praehirsuta*). We used SNP loci that were found to be polymorphic in *J. albifrons* and/or *J. praehirsuta* but monomorphic in the out-group species *J. ischiosetosa* (*n*=9 individuals) and used that information to determine the most likely ancestral allelic state of each SNP. Finally, a single SNP per RAD was randomly retained and this dataset (Brittany: 1913 SNPs, Normandy West: 1082) was projected down to 20 chromosomes per population to optimize JAFS resolution. The JAFS expected from the 13 scenarios were calculated 15 times each and their fit to the observed data was ranked using Akaike information criterion (AIC). The run providing the lowest AIC for each model was kept for comparison between models and estimation of demographic parameters.

### Linkage maps

We estimated recombination between SNP loci using two *J. albifrons* and two *J. praehirsuta* families (supplementary material Table S4). Each family was composed by one male parent and one female parent, and 57 to 70 offspring (full-sibs within each family). The eight parents were born in the lab from controlled crosses, so that each parent was identified and virgin before they were put together by pairs and kept until several broods were produced by each female. Because the average brood size was only ca. 8 offspring, we started with a large number of crosses and kept only the four largest families that we could obtain before the death of the females. All offspring produced were reared individually until they could be sexed and males could be identified. The families used here for building linkage maps were part of the “no-choice” crosses reported in Ribardière et al. (2021), where all experimental conditions are described in detail. Due to a handling error, the mother of one of the *J. praehirsuta* families (family 4, Table S4) could not be genotyped, so that all analyses for this family were conducted with a single parent.

Details of the genotype calling procedure and map construction are given in supplementary material. Briefly, the catalogue of consensus loci in Stacks was built using the seven genotyped parents only, an offspring was conserved only if it had less than 20% missing data, and a SNP locus was conserved only if it had less than 20% missing data in the offspring. The family with the missing mother was treated slightly differently: a locus was kept if at least five copies of the minor allele were present in the family (to discard monomorphic loci). Loci showing significant segregation distortion were filtered out in JoinMap v4.1 (Van Ooijen 2006). Linkage maps were then built using Lep-MAP3 (Rastas 2017) as detailed in supplementary material and visualized using R functions from packages qtl (Broman et al. 2003) and RCircos (Zhang et al. 2013).

Sex chromosomes were identified independently in the three families for which both parents were genotyped. We identified all loci that were heterozygous in at least the mother and had segregation patterns strictly compatible with sex chromosomes and incompatible with autosomes (details in supplementary materials). We compared recombination patterns between sexes on the sex chromosomes by examining the distribution of RAD tags along the corresponding linkage group in male and female maps.

### Genome scan

To look at the variation in genetic differentiation between species across their genome, we used the R package *hierfstat* to calculate *F*_ST_ values for each SNP and plotted them against relative locus positions from one of our linkage maps. We also estimated divergence *d*_XY_ (Nei 1987) for each RAD locus in Stacks. These differentiation and divergence statistics were calculated between *J. albifrons* and *J. praehirsuta* in regions Brittany and Normandy West using the same set of samples used for the demographic analyses performed with *δaδi*. We used the female linkage map obtained from family 1 (*J. albifrons*) as our reference (see supplementary material Table S5A and Fig. S6A). To assess genomic differentiation along the sex chromosomes, we compared the reference female map with a male map (paternal individual from family 1, Table S6), due to observed sex-specific differences in recombination patterns.

Results from the demographic inferences suggested that a large fraction of the genome in *J. albifrons* and *J. praehirsuta* is subject to reduced interspecific gene flow (ca. 60% of the loci in Brittany and 30% in Normandy, see results). To identify which loci were most likely affected by this reduction in gene flow, we used simulations and singled out loci with observed *F*_ST_values apparently incompatible with predictions from a neutral model of evolution (following the logic of Beaumont and Nichols 1996). For that, the software *ms* (Hudson 2002) was used to simulate 200 000 SNP loci evolving neutrally in demographic conditions set to imitate the scenarios inferred previously from *δaδi* analyses. Two simplified scenarios of secondary contact (see results and supplementary material) were thus simulated for regions Brittany and Normandy using parameter values as inferred by *δaδi* for the fraction of the genome that does not involve reduced migration or effective population size (that is, ca. 40% of the loci in Brittany and 70% in Normandy). Then, using our empirical data, we defined the loci most likely affected by gene flow reduction (according to the results from *δaδi*) as those showing a pair of values for *H*_T_(total expected heterozygosity estimated across the two species) and *F*_ST_ that were strictly never obtained by simulation among the 200,000 simulated SNPs (that is, loci with observed *F*_ST_ higher than any simulated SNP for a given *H*_T_).

### Analysis of quantitative trait loci (QTL)

We focused on secondary sexual traits that are known to have a direct role in sexual isolation between species of the *Jaera albifrons* group (Fig. 1). These are specialised spines, lobes and setae distributed on the peraeopods P1-P4 and P6-P7 and used for tactile courtship by males (Bocquet 1953; Solignac 1981; Ribardière et al. 2021). To identify and map the loci potentially linked to the genes encoding these traits, we used an independent dataset obtained from a collection of 15 backcross families designed to show high phenotypic variation in male courtship traits. These families consisted of 3 to 22 full sibs (150 in total), reared under identical control conditions after their production by a cross between an F1 hybrid and either a *J. albifrons* or *J. praehirsuta* (details in supplementary material). We examined the sexual phenotype of each male parent and offspring, recording the presence/absence of a carpal lobe on peraeopods P6 and P7, the number of sexual setae on P1-P4 and P6-P7, and the number of spines on P6-P7. Because the number of setae and spines correlates with the total length of individuals (Prunus 1968), these values were regressed against individual length in generalized linear models, and the standardized residuals of these models were used in QTL analyses. The number of setae had zero-inflated distributions that could not be satisfactorily corrected. Therefore, for P1-4 and P6-7, we considered the presence/absence of sexual setae and then the number of setae when present (details of analysis in supplementary material).

A consensus average genetic map for our 15 backcross families was constructed using Lep-MAP3 as described above. We then used functions from the R packages qtl (Broman et al. 2003) and qtl2 (Broman et al. 2019) to identify QTLs, first by interval mapping and second by genome scan with a linear mixed model (LMM), accounting for relationships between individuals using a random polygenic effect (details in supplementary material). Logarithm of the odds (LOD) significances were tested with 1000 permutation tests. The percentage of variance explained by a QTL was assessed by analysis of variance using type III sums of squares, and confidence intervals were calculated as Bayesian credible intervals. Finally, marker regression (linear regression of phenotypes on marker genotypes) was also used to obtain basic information on the possible presence of QTL, with individuals with missing genotypes discarded.

The genomic regions significantly associated with sexual traits (see results) were reported on our reference linkage map by identifying the RAD-loci at the significant QTL positions and, in return, finding the position of these RAD-loci on our reference map (details in supplementary material).

## Results

### Rate of genotyping error

We found that 0.17% to 0.86% of the locus-by-locus genotypes differed between replicates of a unique individual (using 52997 SNPs). The error rate varied between 0.15% and 0.94% when using a single SNP per RAD-tag (9718 SNPs).

### Genetic structure

The PCA built using all individuals genotyped at 9718 SNPs (supplementary material Fig S2) confirmed the greater divergence of species *J. ischiosetosa* compared to the *J. albifrons* / *J. praehirsuta* pair, as had been reported previously using nuclear AFLP data by Mifsud (2011). This is shown by axis 1 clearly separating *J. ischiosetosa* from the other individuals (5.98% variance explained, Fig. S2). Figure 2B shows the result of a PCA excluding *J. ischiosetosa*, where the first principal component (PC1, 5.76% variance explained) was driven by differences between our two focal species *J. albifrons* and *J. praehirsuta*, and PC2 (4.91%) reflected the spatial distribution of samples. PC3 (3.13%) and PC4 (2.58%) further differentiated samples of different geographic origins (sites Normandy West vs East, and Brittany vs the Isles of Scilly, data not shown). A few individuals characterized by their intermediate phenotype (shown as stars in Fig. 2B) fell in between *J. albifrons* and *J. praehirsuta* on PC1, but most co-occurred with the individuals that had a phenotype typical of either *J. albifrons* (as in site 6, shown in orange in Fig. 2B) or *J. praehirsuta* (site 5, in light blue).

The distribution of the samples along PC1 suggested that mixed *J. albifrons* / *J. praehirsuta* populations are characterized by different levels of genetic differentiation. With the same set of loci used for the PCA, between-species *F*_ ST_estimated in four locations were 0.34 in the Isles of Scilly (location 7 in figure 2A), 0.29 in Brittany (location 1), 0.20 in Normandy West (location 5), and 0.10 in Normandy East (location 6). These differentiation values were all significant (*p* < 0.001).

When we forced TreeMix to represent the history of these four populations as a bifurcating tree without admixture (i.e. no migration allowed between lineages), we obtained inconsistent results. Reciprocal monophyly for *J. albifrons* and *J. praehirsuta* was found in 16 out of 30 trees (Fig. S3A), while in other trees with equal likelihood the position of the Normandy East *J. praehirsuta* population was inconsistent. The residuals of the prevailing tree (reciprocal monophyly, Fig. S3B) suggested that a better fit is expected when allowing for two migration events between species in regions Normandy West and East. With two migration events allowed, 26 out of 30 trees resulted in reciprocal monophyly for the two species with gene flow from *J. albifrons* to *J. praehirsuta* in Normandy West, and in the opposite direction (and stronger) in Normandy East (Fig. 2C). Adding more migration events increased the likelihood of models but added mixtures almost only between Normandy West and East locations (within and between species, data not shown).

### Demographic history of differentiation

Out of thirteen models, the scenario that best explained the observed JAFS for Brittany (Fig. 2D, site 1) was that of a secondary contact with two classes of effective population size and two classes of migration rates (SC2N2m). Secondary contact with two classes of migration rates was also the most likely scenario for Normandy West (Fig. 2D, site 5), but with a single class of effective population size (SC2m). Parameter estimates for the simulation that produced the best fit are reported in supplementary material Table S3. In Brittany, the divergence in allopatry (*t_S_*) preceding the period of secondary gene flow (*t_S_*) accounted for *t_S_*(*t_S_* + *t_SC_*) ≅ 83% of total divergence time, while it amounted to 89% in region Normandy. The fraction of the genome with reduced interspecies migration (1-*P*) was 62% in Brittany and 33% in Normandy, and this reduction effect was on average 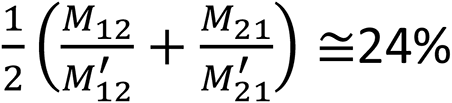 in Brittany and 18% in Normandy. Finally, in region Brittany, a proportion *Q* = 50% of the genome had reduced effective population (reduction factor *H*_rf_ = 0.13).

### Linkage maps

The linkage maps obtained for each of the seven parents contained from 1512 to 4939 SNPs clustered into 10 to 12 linkage groups (Tables S4-11, Fig. S6). The female-heterogametic system of sex determination identified from karyotypes by Staiger and Bocquet (1954) was confirmed here for both species (Table S12). Although both *J. albifrons* and *J. praehirsuta* (like all European species of the *Jaera albifrons* complex) exhibit a large metacentric Z chromosome and two smaller acrocentric W1 and W2 chromosomes, W1 and W2 always segregate together and thus appear as a unique linkage group. Sex chromosomes corresponded to a unique LG common to all individuals in the two species (hereafter labelled LG1).

Figure 3 shows synteny and collinearity between linkage maps for four individuals: a male and a female of each species (parents of the families numbered 1 and 3 in Tables S4-10). These map comparisons based on 1256 to 2182 RAD locus in common between maps suggest intraspecific and interspecific chromosomal polymorphisms. Within species (Fig. 3A-B), LG5 was involved in a potential Robertsonian fusion-scission involving three chromosome arms (called LG5a-c). Comparing maps between species (Fig. 3C-D) revealed an additional fusion-scission (LG6) and a reciprocal translocation (LG8 and 9). These intra- and interspecific chromosomal rearrangements were confirmed when systematically comparing the seven maps two-by-two (Figs. S6 and S7). In particular, LG6 was fused in all four *J. albifrons* maps and cleaved in all three *J. praehirsuta* maps, and the reciprocal translocation in LG8-9 was also confirmed in all interspecific comparisons. The three groups noted LG5a,b,c appeared sometimes to be organized as a-b and c (in *J. albifrons*) or a and b-c (in *J. praehirsuta*), or as three independent groups (in both species). Finally, the *J. praehirsuta* male of family 4, although based on the lowest number of markers (1512, Table S11), confirmed these patterns and showed an additional potential scission involving LG7 (Figs. S6D and S7). The organization of linkage groups of the seven maps is summarized in Figure 3E.

**Figure 3 –.**
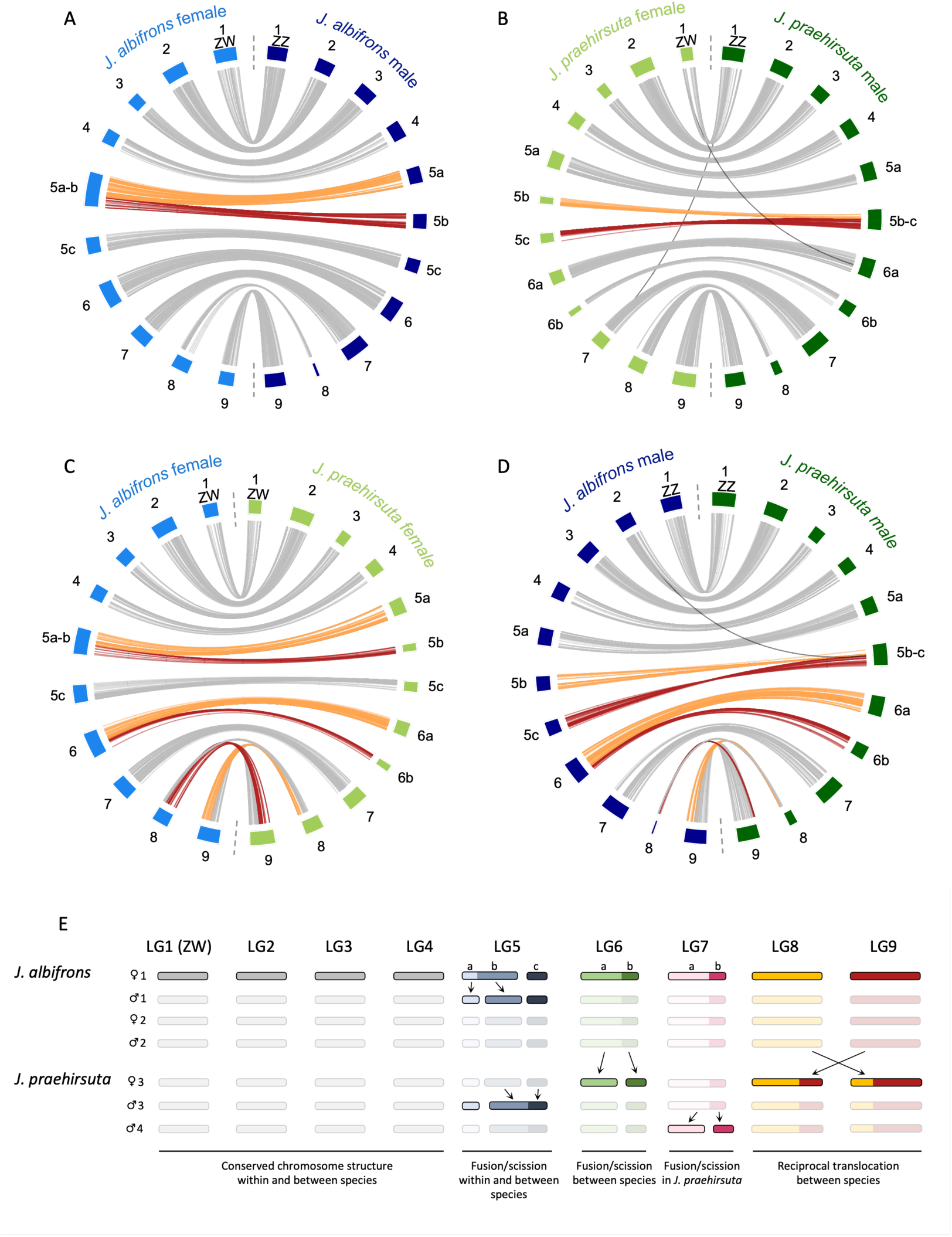
Panels A-D: genome synteny and collinearity between two *J. albifrons* and two *J. praehirsuta* linkage maps (based on 1256 to 2182 SNPs in common between any two maps). Numbered rectangles correspond to linkage groups (LGs). The LGs corresponding to the sex chromosomes were given the number 1. Each SNP common to two maps is shown in grey when it occurs on the same LG in both maps presented on a given chart, and in orange or red when it occurs on different LGs. Putative mapping errors are shown in black. Upper figures: comparison of linkage maps for a male and a female of the species *J. albifrons* (A) or *J. praehirsuta* (B). It can be seen that linkage groups 5a-c are involved in putative fusion/scission chromosome rearrangements within each species. Bottom: the same data are used to compare males (C) or females (D) from each species. These species comparisons reveal an additional potential fusion/scission (LG 6) and a reciprocal translocation (LGs 8-9). The maps used in this figure are those that were obtained for the parents of family 1 (*J. albifrons*) and 3 (*J. praehirsuta*). Panel E shows a summary of putative chromosomal polymorphism inferred from the seven linkage maps obtained in this study. The size of each linkage group has been arbitrarily set for illustrative purposes only, but colours indicate synteny between linkage groups across individuals, and a-c letters are as in Supplementary Fig. S6. Grey linkage groups show no evidence of rearrangements across all individuals, while other colours help to visualise chromosomal rearrangements compatible with Robertsonian fusion/scissions (LG5-7) and a reciprocal translocation (LG8-9).

### Genome scan and QTL

Figure 4 shows the distribution of genetic differentiation between species (measured as *F_ST_*_,_ at each SNP) along a linkage map used as reference (*J. albifrons* female, family 1, Fig. 3A). In the two regions considered in this analysis (Brittany and Normandy West), locus-specific *F_ST_*, varied from 0 to 1, and a large number of SNPs had stronger differentiation than that of any simulated data (2694 out of 35442 SNPs = 7.6% in Brittany, 964 out of 45840 = 2.1% in Normandy West, shown in red in Fig. 4 for those that were also present on the reference map). In Brittany, there were regions of very strong differentiation on all linkage groups, and the distribution of differentiation within each of these groups was heterogeneous. By contrast, in region Normandy, the distribution of genetic differentiation was more variable amongst linkage groups. In particular, differentiation was much reduced in LGs 2, 3, 5c, and to a lesser extent 7, than in the other ones. Sharp "peaks" of differentiation were visible on LG1 (sex chromosomes), and on the LG involved in chromosomal rearrangements between species (LG5a-b, LG6, LG8-9). A region of strong differentiation was also visible on LG4. Divergence, measured as *d*_XY_, show similar patterns (Fig. S10).

**Figure 4 –.**
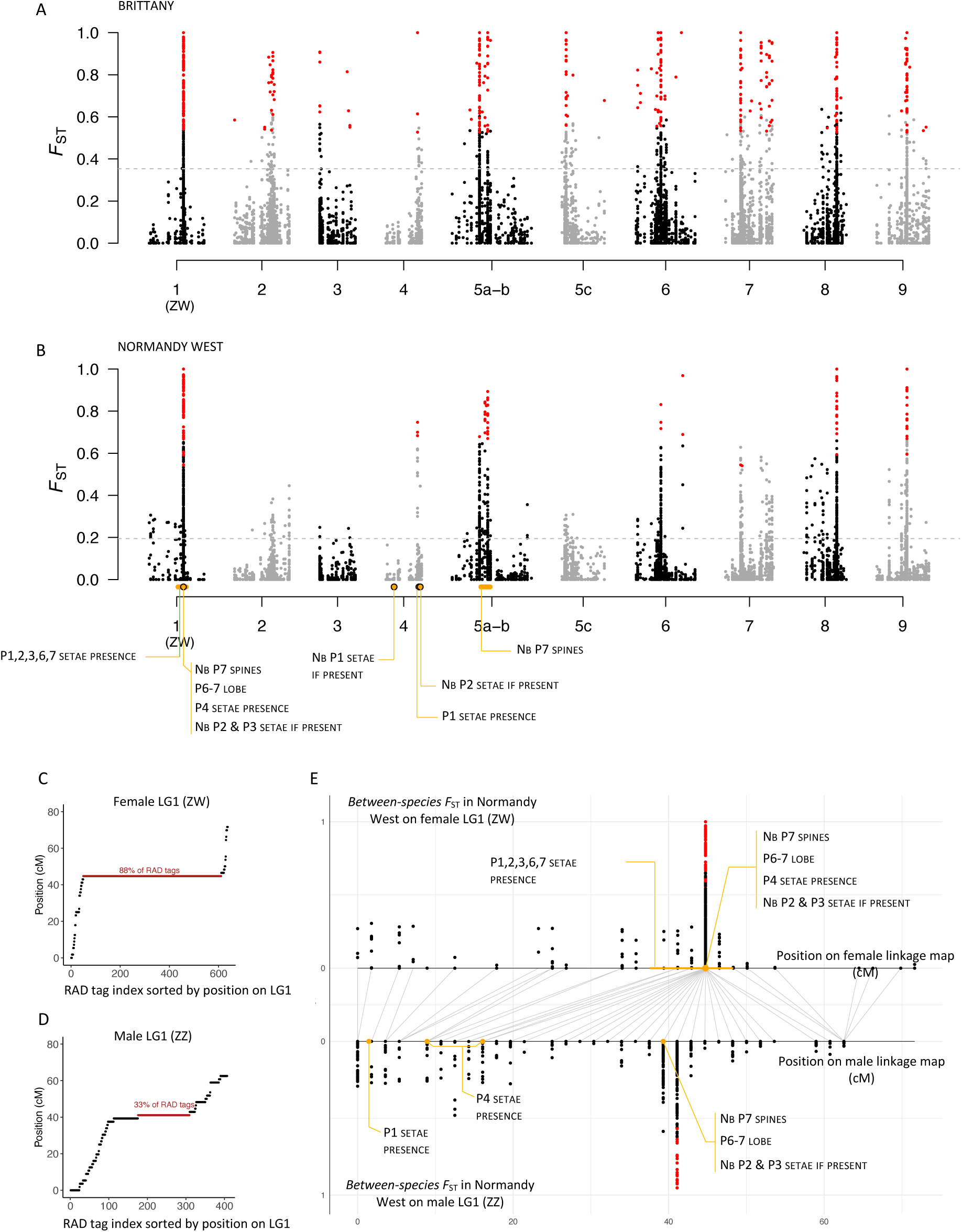
Distribution of genetic differentiation (*F*_ST_) between *J. albifrons* and *J. praehirsuta* in Brittany (panel A: strong ecological and sexual isolation, reduced introgression) and Normandy West (B: no ecological isolation, slightly reduced sexual isolation, introgressive hybridization in progress). Each point represents the *F*_ST_ value between species measured at one SNP (here 9028 and 9292 SNPs, respectively). The reference map used here is that of a female *J. albifrons* (maternal map of family 1, shown in light blue in Fig. 3A). Observed *F*_ST_ values that were greater than any value obtained by simulating 200 000 SNPs evolving with genome-wide homogeneous gene flow are shown in red. The horizontal dashed lines give the genome-wide *F_ST_* (i.e. calculated over all SNPs) in each region. Significant QTL loci are represented here by an orange dot or segment depending on the accuracy with which the QTL locus could be mapped on the reference linkage map used here. In Normandy, the action of gene flow between species has resulted in a highly heterogeneous erosion of genetic differentiation, leaving strong differentiation on sex chromosomes (LG 1), rearranged chromosomes (LGs 5, 6, 8, 9), and chromosome 4. Panels C-E focus on sex chromosomes. Panels C and D show the distribution of RAD tags on linkage group 1 (sex chromosomes) in a female (panel C, reference female map) and a male (D, *J. albifrons* father in family 1 used here as an example, see Fig. S8 for similar representations for all linkage maps). RAD sequences are sorted according to their position on LG1 (in cM). A region of low or null recombination can be identified from the concentration of markers (in red: single position aggregating the largest number of RAD-tags). Panel E shows the distribution of genetic differentiation (*F*_ST_) between *J. albifrons* and *J. praehirsuta* in the Normandy West population, where introgressive hybridization is ongoing. Each point represents the *F*_ST_ value at one SNP plotted on the female map (above zero) and the male map (below zero). Grey lines connect RAD tags shared between the male and female maps, highlighting differences in recombination patterns between ZW and ZZ chromosomes.

Interval mapping analyses (Table S15) showed that 11 male secondary sexual traits (out of 16 analysed) were significantly associated with a 14.5 cM region on linkage group 1 (sex chromosomes, Fig. 4). Three of these traits were also significantly associated with genetic variation at three other positions on LG 4 and 5a-b, and one additional trait was associated with a distinct position on LG 4 (Figure 4). In total, therefore, twelve traits were associated with a few (possibly physically very large) genomic regions located on three linkage groups.

All traits but one (number of setae on P3) were also identified by the Genome Scan LMM method. These two methods agreed on the most likely location of all characters, except for the number of setae on P2, which was located on the same LG but not at the same position. Finally, the less powerful locus-by-locus regression method found a single locus on LG1 to be significantly associated with 9 traits, confirming the results of the interval mapping and LMM methods for these traits.

The distribution of RAD tags on the linkage group corresponding to the sex chromosomes reveals a substantial region of recombination suppression. This suppression is more pronounced between the Z and W chromosomes, with 81 to 88% of RAD tags mapping to a unique position on female linkage maps (Fig. 4C, S8A–C). A similar, though less extensive, pattern is observed between Z chromosomes, where 23 to 58% of RAD tags are localized to a single position in the center of the linkage group (Fig. 4D, S8D–F). By comparing the distribution of genetic differentiation and sexual phenotype QTLs along a female and a male linkage map (parents of *J. albifrons* family 1), Figure 4E shows that five of the seven QTLs that could be located on the male map were found within a central region of the Z chromosome that does not recombine in either sex.

## Discussion

### Secondary contact has resulted in different levels of interspecific gene flow between J. albifrons and J. praehirsuta

Phenotypic observations (Solignac 1969, 1978) and microsatellite data (Ribardière et al. 2017) led to the conclusion that *J. albifrons* and *J. praehirsuta* experience introgressive hybridization in one region of the French coast (Normandy) while they are strongly isolated in other areas (e.g. Brittany). This contrast is essential for interpreting the genome-wide distribution of genetic differentiation in terms of gene flow, while excluding confounding factors such as local reduction in nucleotide diversity due to genetic hitchhiking and background selection (Noor and Bennett 2009). Therefore, our first aim was to use genome-wide SNP data to refine these findings and infer historical scenarios of population divergence.

The distribution of genetic variation observed at nearly 10 000 SNPs (roughly 0.001% of the genome considering all variable and non-variable sites) confirmed that interspecific differentiation varies with geography. The principal component analysis of multilocus genotypes (Fig. 2B) showed a gradient from strong differentiation (Isles of Scilly, *F*_ST_= 0.33) to mild differentiation (Normandy East, *F*_ST_=0.10), and the best-fit tree inferred by Treemix required two migration events between *J. albifrons* and *J. praehirsuta* in the two geographic locations with lower differentiation (Fig. 2C). Demographic inference using δaδi indicated that in both high (Brittany) and low (Normandy) gene flow regions, a secondary contact model best explained the data, with 83–89% of total divergence time spent in allopatry (Table S3). Together with the PCA, *F*_ST_ and Treemix results, these findings support the conclusion that re-contact after isolation has resulted in markedly different levels of interspecies gene flow in different areas where the species met.

These results are consistent with Bocquet and Solignac’s conclusion that hybridization is "exceptional" in Brittany (based on extensive phenotypic observations, Bocquet and Solignac 1969) rather than totally absent (Ribardière et al. 2017). In this region, where the two species occupy distinct intertidal habitats, ecological isolation was estimated to 98% (Ribardière et al. 2021), and complemented by strong sexual isolation (estimated to 73% under no-choice conditions if this barrier was acting alone). These two barriers together amount to 99% reproductive isolation. It is thus possible that rare hybridization events occur when individuals meet on the same substrate and bypass sexual barriers, a process that could be captured by the analysis of joint allelic frequency spectra.

In comparison, in Normandy, where suitable sheltered algal habitats are not available, reproductive isolation is maintained by behavioural sexual barriers alone (estimated to 37% and 92% using no-choice and free-choice experiments, respectively, Ribardière et al. 2021) Our genetic results support the view that this incomplete isolation allows for introgressive hybridization (Bocquet and Solignac 1969; Ribardière et al. 2017, 2021). Interestingly, in the Isles of Scilly (site 7 in Fig. 2A), genetic differentiation between the two species is high despite the apparent absence of ecological isolation, suggesting that stronger sexual isolation (or other unidentified barriers) may be sufficient to prevent gene flow in this region. This confirms the sharp contrast in heterospecific gene flow between geographic regions, providing an interesting opportunity to investigate how differences in reproductive isolation shape genomic differentiation between *J. albifrons* and *J. praehirsuta*.

### The genomes of Jaera albifrons and J. praehirsuta are semi-permeable to gene flow

Genetic differentiation (*F*_ST_) along linkage groups varied widely in both geographic areas (Fig. 4), including several genomic regions with very high differentiation (up to complete fixation of alleles between species, *F*_ST_ = 1). However, if one were to look at each of the two differentiation landscapes shown in Figure 4 independently, it would remain difficult to determine whether these patterns are caused by variations in gene flow along the genome.

In Brittany, where there are strong peaks of differentiation across all linkage groups, the effect of barrier loci (including habitat isolation) cannot be distinguished from accumulated differentiation and divergence genome-wide due to background selection and drift (reviewed in Ravinet et al. 2017). In Normandy, where differentiation peaks are more localized (Fig. 4B), heterogeneity could also originates from local variations in nucleotide diversity due to selective sweeps and background selection rather than differences in gene flow (Noor and Bennett 2009; Roesti et al. 2012; Cruickshank and Hahn 2014; Seehausen et al. 2014; Burri et al. 2015; Ravinet et al. 2017; Sætre and Ravinet 2019). This is especially relevant in regions of low-recombination, such as those associated with centromeres (Noor and Bennett 2009; Talbert and Henikoff 2010), chromosomal rearrangements (Noor and Bennett 2009), and sex chromosomes (Presgraves 2018; Van Belleghem et al. 2018). Here, all three factors may account for some of the heterogeneity observed in the genetic differentiation between species. Cytological observations for populations close to our sampling sites in Brittany have shown i) a majority of metacentric chromosomes and a few acrocentric ones in both species (Lécher 1967b), ii) intra- and interspecific chromosome number polymorphism attributed to Robertsonian fusion-scissions (Staiger and Bocquet 1956; Lécher 1967b; Lécher and Prunus 1971), and iii) female-heterogametic sex determination with a large metacentric Z chromosome and two smaller acrocentric W chromosomes (Staiger and Bocquet 1954; Lécher 1967b). Our linkage maps are consistent with these previous observations, and the density of markers along these maps suggest that there are large low- or no-recombination regions where differentiation peaks could reflect local linked selection rather than reduced interspecies gene flow. This mechanism is further supported by within-species comparisons across geographic regions, where differentiation mirrors that observed between species in Brittany (Supplementary Fig. S11). Such patterns driven by geographic isolation and habitat fragmentation rather than reproductive isolation (Ribardière et al. 2021) reinforce the idea that local genomic differentiation is shaped by factors beyond interspecies semi-permeability to gene flow.

However, the semi-permeability of these genomes to gene flow is revealed by comparing differentiation patterns between the two regions studied, where we have shown that interspecies gene flow is highly unequal. Following the idea of comparing the differentiation landscapes observed in different geographic areas (Noor and Bennett 2009; Harrison and Larson 2016; Le Moan et al. 2016; Wolf and Ellegren 2017; Sætre and Ravinet 2019; Westram et al. 2021), we identified genomic regions maintaining strong differentiation despite gene flow. As predicted by Wu (2001; see also e.g. Feder et al. 2012), some high-differentiation regions in Brittany showed no or little homogenizing effect of introgression in Normandy’s hybridizing populations. Specifically, six genomic regions on linkage groups 4-6, 8-9, and the sex chromosomes (LG1), resist introgression in Normandy, while differentiation elsewhere is markedly eroded (with *d*_XY_ patterns showing broad similarity, Fig. S10). Regardless of local variations in recombination rate, the random segregation of chromosomes across generations should homogenize across species the allelic frequencies not linked with species-defining sexual phenotypes.

In line with these results, nearly all sexual QTLs appeared to be in strong linkage (same position on linkage maps) with the highly differentiated regions that resist introgression (Fig. 4). Three regions with strong differentiation peaks (on LG 1, 4, and 5) coincided with the presence/absence or number of sexual setae, lobes, and spines used for courtship. Only one sexual trait (number of setae on P1) was found to be associated with genetic variation at a position of the linkage map where the two species show little differentiation (LG4, and that is true in the two geographic regions analysed). We stress that these observations are based on recombination maps and not physical genomic positions, so that here the "coincidence" of sexual QTLs and differentiation peaks indicates genetic linkage across potentially large intervals.

We conclude from these observations that gene flow has eroded differentiation unevenly across the genome, and that at least three regions resisting this erosion harbour genes directly involved in behavioural sexual isolation.

Additionally, three other differentiation hotspots (on LG 6 and 8-9, Fig. 4) were not associated with any observed sexual traits. Given their strong genetic differentiation (up to complete fixation of distinct alleles in the two species, in sharp contrast with the reduced differentiation in other linkage groups), these regions could also contain barrier loci involved in reproductive isolation yet to be identified. Such loci might govern sexual isolation through unobserved traits (e.g. chemical signals, female preferences) or other isolating barriers (e.g. genetic incompatibilities).

Whether or not associated with identified barrier effects, these regions impermeable to gene flow were not randomly distributed, concentrating on sex chromosomes and rearranged chromosomes (Fig. 4).

### Sex chromosomes carry a genomic region that is resistant to interspecies gene flow

Sex determination was established by Staiger and Bocquet (1954, 1956) for the four European species of the *Jaera albifrons* complex. A trivalent element observed exclusively in females at meiotic metaphase allowed these authors to conclude that sex determination involved a ZW_1_W_2_ / ZZ system, and that this system most likely evolved from a Robertsonian fusion between ancestral ZW sex chromosomes and a pair of autosomes (it was also the first identification of sex chromosomes for any isopod, Vandel 1947; Staiger and Bocquet 1954). Here, we could confirm that females are the heterogametic sex, and that *J. albifrons* and *J. praehirsuta* share the same pair of sex chromosomes (noted LG1 in our analyses). Linkage maps built from observations of recombination events are unable, however, to distinguish W_1_ and W_2_ chromosomes since these two are always segregating jointly (hence ZW_1_W_2_ will appear as a single linkage group, as in Figs. 3-4 and S6-7).

Interestingly, sex chromosomes showed the sharpest differentiation peak between species and concentrated most of the genetic variation found to be associated with male sexual traits (11 traits, Fig. 4).

We identified 274 to 335 sex-linked RAD loci mapping to a single position within each female recombination map (corresponding to "position 44.7 cM" on our female reference linkage map, Fig. 4, and supplementary figure S8 and Table S15). These loci were thus all located in a sex-linked region of unknown physical size harbouring the master sex-determining gene and where Z and W chromosomes do not recombine within species. Interestingly, we also observed that recombination is suppressed in the central region of the Z chromosome in males, as evidenced by the distribution of RAD sequences on linkage group 1 (Fig. 4D, supplementary Fig. S8). Suppressed recombination in the central Z region possibly reflects reduced recombination near the centromere of this large metacentric chromosome thought to result from the fusion of two acrocentric chromosomes (Staiger and Bocquet 1954).

Looking at the genetic differentiation between species in the hybridizing populations of Normandy West (Fig. 4E), we saw that this sex-linked region aggregated the vast majority of SNPs for which a *F*_ST_ was obtained (942 out of 1024 SNPs, i.e. 92%), including all 117 outliers that had *F*_ST_ value higher than that of any of 200 000 simulations with unrestricted interspecies gene flow. A similar pattern was observed when using a male reference linkage map: for example, using the *J. albifrons* male from family 1, 203 out of 818 SNPs with an *F*_ST_ estimate mapped to a single position corresponding to the non-recombining region of the Z chromosome, including all 37 *F*_ST_ outliers (Fig. 4E).

Six male sexual QTLs were associated with the sex-linked genomic region (position 44.7cM on the reference female map, Fig. 4E), and the five remaining QTLs found on the sex chromosomes were located in a tightly linked genomic region (37.6 to 48.3 cM). Using a male linkage map as reference, five of the sexual phenotype QTLs were located within the non-recombining region of the Z chromosome, while two others could be mapped to a region where ZZ recombination occurs (Fig. 4E).

Our findings are consistent with other studies reporting sex-linked QTLs associated with sexual traits (Dopman et al. 2004; Kitano et al. 2009; Lagisz et al. 2012; Liu and Karrenberg 2018; Berdan et al. 2020). In particular, they align with empirical observations suggesting a strong Z chromosome effect in sexual isolation (Qvarnstrom and Bailey 2009). Sex chromosomes could be enriched with sexual trait genes because linkage with the sex-determination locus could facilitate the evolution of sexual traits towards sex-specific, divergent optimal values (e.g. Irwin 2018). However, this idea is debated, as it has received mixed empirical support (e.g. Lande 2000; Beukeboom and Perrin 2014). Moreover, the conflicts generated by divergent sex-specific optimal trait values can also be solved by the sex-specific expression of autosomal genes (a pattern with more empirical support). It is important to interpret these results cautiously. In our case, QTL detection was likely impacted by suppressed recombination in a possibly large region of the sex chromosomes (Noor et al. 2001a; Noor and Bennett 2009; Roesti et al. 2012; Koch et al. 2021). This limits the resolution of our analysis, preventing us from precisely identifying the number of genes involved and their exact genomic locations.

Nevertheless, our results point to the presence of an extensive non-recombining region on the Z chromosome that appears to i) harbour an unknown number of genes that contribute to the determinism of male sexual traits used in courtship, and ii) resist introgression in the face of ongoing hybridisation. Further work using high-quality reference genomes is required to identify these genes and determine whether they are linked to loci controlling female preference, key steps toward understanding the evolution of this genomic architecture.

Beyond the co-evolution of traits and preferences involved in sexual isolation, the asymmetric inheritance and unique recombination patterns of sex chromosomes can facilitate the evolution of other reproductive barriers, especially genetic incompatibilities (reviewed in Qvarnstrom and Bailey 2009; Beukeboom and Perrin 2014; Seehausen et al. 2014; Irwin 2018; O’Neill and O’Neill 2018; Payseur et al. 2018). In flycatchers and Gouldian finches, for example, the Z chromosome harbours barrier loci involved in male traits and female preferences, but also genetic incompatibilities (Saetre et al. 2003; Saether et al. 2007; Pryke and Griffith 2009; Pryke 2010; see also Kitano et al. 2009 for an example involing the X chromosome in sticklebacks). In *J. albifrons* and *J. praehirsuta*, however, first-generation post-zygotic isolation appears minimal: F1 hybrids exhibit normal survival and fertility (Ribardière et al. 2021) and hybrid males were fertile enough to produce 150 offspring from 15 backcrosses used in our QTL analysis. These observations contrast with the more divergent *J. albifrons* / *J. ischiosetosa* pair, where sex-linked post-zygotic barriers reduce hybrid viability (reduction in viability and fertility of F1 hybrids, Solignac 1976). Nevertheless, we currently lack detailed fitness data for backcross and F2 generations, which are required to more accurately quantify post-zygotic isolation and assess whether any such effects are sex-linked.

Finally, while we have obtained a useful description of sex chromosome recombination within each species, we lack a description of the recombination landscape as it would unfold during meiosis in hybrids. This could be obtained by building linkage maps for F1 hybrid parents, which would allow us to estimate ZZ and ZW recombination when the sex chromosomes come from different species. In combination with estimates of post-zygotic barrier effects, this information will be necessary to assess if sex chromosomes promote reproductive isolation through fitness effects of genetic incompatibilities and/or recombination arrest impeding the mixing of alleles involved in male signalling and perhaps other barrier loci (e.g. Rosser et al. 2022).

The same reasoning and perspectives apply to chromosomal rearrangements, which, besides sex chromosomes, appear to play a major role in the reproductive isolation between our two *Jaera* species.

### Rearranged chromosomes play another key role in reproductive isolation

Four of the five strongly differentiated genomic regions (outside sex chromosomes) were located on linkage groups that show structural variation between species: LG5 seems to be composed of three chromosomal arms (here noted a-c) that show different Robertsonian arrangements within and between species, LG6 is divided into two groups in *J. praehirsuta*, and LG8-9 show a reciprocal translocation between species. While the translocation had not been detected before, our results regarding Robertsonian fusions / fissions agree with the chromosomal polymorphism (intra- and inter-species) previously described through karyotype studies for *J. albifrons* and *J. praehirsuta* (Staiger and Bocquet 1956; Lécher 1967b; Lécher and Prunus 1971).

There was in fact only one exception to high genetic differentiation being associated either with sex or rearranged chromosomes: LG4 showed a genomic region resistant to gene flow without any obvious evidence of structural variation between species. As mentioned above, this linkage group harbours genetic variation associated with three male sexual traits characterizing the number of courtship setae on anterior pereiopods. The high density of markers in this genomic region is consistent with reduced recombination, possibly due to proximity to the centromere on an acrocentric chromosome. If suppressed recombination extends to regions that include coding genes near the centromeric region, it could limit genetic mixing and help preserve male sexual trait variation from being eroded by gene flow. Alternatively, given that our linkage maps were based on modest numbers of offspring (which limits the detection of recombination, and thus locus ordering), we also cannot refute the possibility of an undetected, small scale, rearrangement (such as a relatively small inversion, for instance). Careful examination of the collinearity of the markers on LG4 (Fig. S6) showed no evidence of such a rearrangement, but it cannot be ruled out that denser genetic maps or genome sequences will reveal new structural variants.

Like sex chromosomes, structural variants can promote reproductive isolation through two mechanisms. First, they can have direct detrimental fitness effects in hybrids that are heterozygote for the rearrangements (White 1973). This occurs when pairing, chiasma formation, or segregation do not proceed properly at meiosis, resulting in germ cell death or unbalanced (aneuploid) gametes, which reduce the fertility of hybrids (White 1978; Searle 1993; reviewed in Faria and Navarro 2010). Fusions/ fissions or translocations such as observed in this study can potentially have such direct effects on the fitness of heterokaryotic hybrids (reviewed in Berdan et al. 2024). The idea of a causal relationship between Robertsonian polymorphism and speciation in isopods can actually be traced back to Vandel (1947).

However, the underdominance of individuals heterozygous for such rearrangements is far from automatic (e.g. Rieseberg 2001), and two sets of observations even suggest the opposite in the case of *J. albifrons* and *J. praehirsuta*. First, as explained above in the context of sex chromosomes, preliminary observations rather point towards no reduction in F1 hybrid viability, and no or limited reduction in their fertility. Second, at least in the case of LG5, the organization of the three chromosomal arms noted a, b, and c in Fig. 3 seems to be irrelevant to (direct) reproductive isolation since LG5 shows structural polymorphism within each species. This structural variation is fully compatible with the variation in chromosome number described between populations within *J. albifrons*. The chromosomal polymorphism of this species was studied extensively, revealing an impressive variation in the number of chromosomes, ranging from 9 to 14 pairs along Europe’s coasts (Staiger and Bocquet 1956; Lécher 1964, 1968; Lécher and Prunus 1971). Analyses of chromosome arm length, centromere distribution, and nuclear DNA content showed that this polymorphism was compatible with Robertsonian rearrangements (Lécher 1967a,b) and did not cause reproductive isolation (Lécher 1964, 1967b). In fact, despite being the most prominent advocate of the subdominant chromosomal rearrangement theory, M. J. D. White himself questioned the impact of chromosomal rearrangements on fitness in the *Jaera albifrons* complex (White 1978).

The reciprocal translocation observed here (LG 8-9 in Fig. 3) is a new finding, but even this type of rearrangement does not automatically lead to reproductive isolation. During meiosis, the four chromosomes involved can form a quadrivalent structure, followed by the production of a variable proportion of viable gametes, depending on how the chromosomes segregate.

The second class of effect of chromosomal rearrangements is through recombination suppression (suppressed-recombination models, Trickett and Butlin 1994; Noor et al. 2001b; Rieseberg 2001; Faria and Navarro 2010). Recombination is suppressed near fusion-scission and translocation breakpoints, which could allow barrier loci to become protected from introgression and associations of alleles to become associated. The co-occurrence of a QTL for carpal spine number and strong interspecies genetic differentiation near the breakpoints between LG5a and b (Fig. 4) strongly supports this hypothesis, but as with the sex chromosomes, more information is needed on the genetic determinism of sexual traits at this site, the association of loci encoding sexual and other types of barriers, and the direct effect of chromosomal rearrangements on post-zygotic segregation.

## Conclusion

The genomic analyses presented here show that the marine isopods *Jaera albifrons* and *J. praehirsuta* experience extremely low heterospecific gene flow in one region of western France (Britany) but strong introgressive hybridisation in another (Normandy), confirming previous conclusions based on observations of phenotypes in natural populations (Solignac 1969), experimental crosses (Bocquet and Solignac 1969), and low-coverage genetics (Ribardière et al. 2017). This contrast allows us to conclude that six genomic regions show stronger interspecies differentiation than expected in Normandy because they resist introgression, while the rest of the genome is much more permeable to gene flow. All but one of the genomic regions resistant to interspecies gene flow are located either on the sex chromosomes or on rearranged chromosomes (several fusion-scissions and one reciprocal translocation). These genomic regions show low recombination and, in two cases, genetic variation associated with several male courtship traits. In particular, the sex chromosomes were found to carry genetic variation associated with the number of sexual setae, lobes and spines, i.e. all aspects of the traits used by males in courtship.

The coincidence of outlier loci and sexual QTL on non-recombining regions suggests that suppression of recombination may have been critical for the evolution and maintenance of reproductive isolation by preventing mixing of genetic material in regions harbouring sexual barrier loci (in agreement with predictions by e.g. Noor and Bennett 2009; Faria and Navarro 2010). Recombination suppression may result from various factors such as proximity to centromeric regions, which can occur on both sex chromosomes and autosomes, as well as breakpoints of major chromosomal rearrangements. However, unequivocal demonstration of this hypothesis still requires estimates of recombination between sex and rearranged chromosomes in heterokaryotic individuals, and formal tests of the direct barrier effect of sex and rearranged chromosomes through estimates of post-zygotic isolation beyond the first generation of hybrids. In particular, the role of fission-fusion and translocations is less well understood than that of inversions (Lucek et al. 2022; Augustijnen et al. 2024). Moreover, we currently have no information on the physical size of the genomic regions involved, which represents a significant limitation. The use of reference genomes will be essential to accurately characterize these regions and advance our understanding of their contribution to reproductive isolation.

Our results also show that barrier loci must be present in at least one other region of the genome (where very strong differentiation was observed outside rearrangements) and that other barrier loci (e.g. genetic incompatibilities, sperm competition, chemical communication) may co-localise on sex chromosomes or rearranged regions (e.g. near translocation breakpoints where no QTL were found). We also need to determine whether the chromosomal differences we have observed are fixed between species, and to obtain more precise information on the genetic determinism of sexual traits. Taken together, our results suggest that a long period of allopatry could have enabled the divergent co-evolution of male traits and female preferences, involving at least some genes located in non-recombining regions on sex chromosomes and rearranged chromosomes. However, the timing of events remains elusive, particularly regarding the order of appearance of premating isolation versus chromosomal rearrangements and sex chromosome divergence, and their respective roles before and after secondary contact.

## Supplementary information

A single PDF file containing all supplementary material is publicly available on Biorxiv together with this article at https://www.biorxiv.org/content/10.1101/2025.01.08.631900v2.supplementary-material

## Data, script and code availability

RAD-seq reads are publicly available on NCBI (bioproject PRJNA1173521, accession numbers SAMN44307079 to SAMN44307994). The other data and scripts associated with this study are available on the zenodo digital repository at https://doi.org/10.5281/zenodo.14214943 (Broquet 2024).

## Author contributions

Conceptualization and methodology: AR, CDT, KA, PAG, and TB. Field sampling and species identification: AR, JC, CH, SL, and TB. Crossing experiments and maintenance of the individuals in the laboratory: AR, JC, and TB. Genotyping and phenotyping: AR, CDT, JC, GLC, CH, and TB. Analyses and writing: AR, KA, PAG, and TB. Supervision, project administration and funding acquisition: TB.

## Funding

This study was funded by the French Agence Nationale de la Recherche (grant ANR-13-JSV7-0001-01 to T.B.).

## Conflict of interest

The authors declare they have no conflict of interest with the content of this article. T. Broquet and P-A. Gagnaire are recommenders for PCI evolutionary biology.

## Acknowledgements

We thank Alan Brelsford for sharing his knowledge on the lab work and bioinformatics of ddRAD-seq. We thank Alan Le Moan, Stuart Piertney, and Marius Wenzel for insightful discussions, and Caroline Broudin and Olivier Timsit for their help with sampling in Normandy. We are grateful to Denis Roze for the sampling mission to the Isles of Scilly aboard *Pen Kreo*. We thank Sébastien Colin for the confocal laser scanning microscope images used in Figure 1. This work benefited from access to the EMBRC-France and Biogenouest Genomer platform at the Station Biologique de Roscoff and the Roscoff Bioinformatics platform ABiMS.

## Notes

https://doi.org/10.5281/zenodo.14214943

### Competing Interest Statement

The authors have declared no competing interest.

### Summary of Updates

PCI badge added following recommendation of this preprint by Peer Community in Evolutionay Biology. Minor text corrections.

