## Supplementary material for "Sex chromosomes and chromosomal rearrangements are key to behavioural sexual isolation in *Jaera albifrons* marine isopods"

and Broquet T.

SAMPLING 2
DOUBLE-DIGEST RESTRICTION ASSOCIATED DNA SEQUENCING (DDRADSEQ) 3
RAD-seq analysis parameters fitted to suit each objective 4
GENETIC STRUCTURE 5
ACP 5
Treemix 6
DEMOGRAPHIC HISTORY OF DIVERGENCE 7
*δ*a*δ*y 7
LINKAGE MAPS 10
IDENTIFICATION OF SEX CHROMOSOMES 13
GENOME SCANS 17
Simulations 17
QTL ANALYSIS 21
Families 21
Phenotypic traits 21
Genetic map 23
Location of QTL on reference linkage map 23
REFERENCES 24

### Sampling

**Table S1**: Localisation of the sampling sites and number of *J. albifrons* and *J. praehirsuta* males used in the analyses.

| Area | Sampling sites | Latitude | Longitude | Phenotypes | | |
|---|---|---|---|---|---|---|
| | | | | *J. albifrons* | Intermediate | *J. praehirsuta* |
| 1 | 1.1 | 48°40'27.59"N | 3°57'11.30"W | 15 | 0 | 10 |
| | 1.2 | 48°40'20.86"N | 3°57'0.32"W | 15 | 0 | 9 |
| | 1.3 | 48°39'10.84"N | 3°57'2.99"W | 20 | 0 | 20 |
| 2 | | 48°50'22.87"N | 1°35'56.35"W | 3 | 0 | 0 |
| 3 | | 49°43'5.35"N | 1°57'0.92"W | 5 | 0 | 1 |
| 4 | | 49°34'54.86"N | 1°15'57.27"W | 8 | 0 | 0 |
| 5 | 5.1 | 49°23'25.26"N | 1° 2'5.04"W | 20 | 2 | 18 |
| | 5.2 | 49°21'17.53"N | 0°47'59.78"W | 20 | 11 | 20 |
| | 5.3 | 49°20'53.84"N | 0°40'55.92"W | 15 | 0 | 1 |
| 6 | | 49°44'24.30"N | 0°18'38.46"E | 45 | 11 | 12 |
| 7 | | 49°53'38.94"N | 6°20'14.16"W | 38 | 1 | 5 |

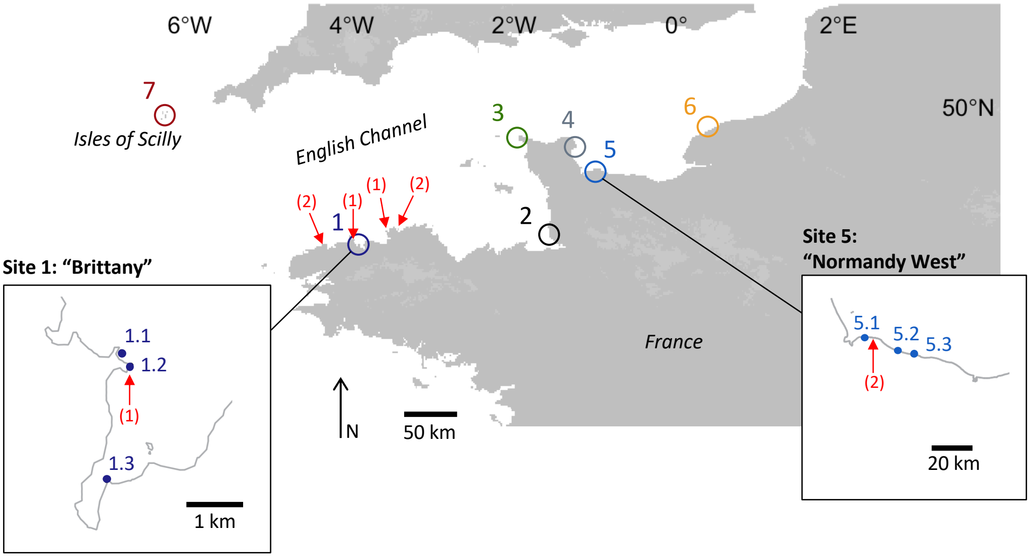


Figure S1 - sampling sites, including *J. ischiosetosa* samples. The analyses that are restricted to region "Brittany" and region "Normandy West" correspond to sampling sites 1 (1.1, 1.2 and 1.3) and 5 (5.1 and 5.2), respectively. The nine *J. ischiosetosa* males used as outgroup came from six sites indicated by the red arrows (with the number of individuals in brackets).

#### Double-digest restriction associated DNA sequencing (ddRADseq)

All genetic analyses were based on bi-allelic SNP genotypes obtained from double-digest RAD sequencing (Peterson et al. 2012). Genomic DNA were extracted using Nucleospin Tissue kits (Macherey–Nagel), and individual libraries were produced from 50ng of DNA using PstI and MseI restriction enzymes following a protocol modified from Brelsford et al. (2016) and detailed in Daguin-Thiébaut et al. (2021). With this protocol, individual libraries are pooled only after amplification, allowing us to homogenize the contribution of each individual to the final pooled library and to the final sequence dataset. A total of 942 samples (NCBI Bioproject PRJNA1173521) were included in 6 different libraries that were sequenced as 125 bp single-end reads using Illumina HiSeq 2500 v4 technology either by Eurofins Genomics (Germany) in 2015 or Fasteris (Switzerland) in 2017). Library preparation and sequencing included 14 pairs of replicates (i.e. all steps of lab work after DNA extraction replicated once for 14 individuals) in order to estimate genotyping error rates.

The quality of raw sequencing data was controlled using *fastqc* (Andrews 2010) and *multiqc* (Ewels et al. 2016). Illumina adapters were removed using *cutadapt* (Martin 2011) and then all reads were trimmed to 110 bp using *trimmomatic* (Bolger et al. 2014). Alignment and snp calling was then performed in Stacks v2.52 (Catchen et al. 2011, 2013) using components of the *denovo_map* pipeline with parameters adjusted to suit the purpose of our different objectives (see supplementary material). The data produced by Stacks were further filtered in *R* v3.5.3 (R Core Team 2019) to remove SNP loci and individuals with more than 10% to 30% of missing data depending on the objective (see below and supplementary material). Finally, the rate of genotyping error was estimated by counting the number of genotypes that differed within each of our 14 replicate pairs (i.e. error rate = number of differences / number of markers genotyped in both replicates of a pair). One replicate of each pair was then randomly removed from the dataset for downstream population genetic analyses.

#### RAD-seq analysis parameters fitted to suit each objective

**Table S2:** summary of analysis parameters, filters based on missing data, and final number of individuals and locus used in dowstream analyses. Error rates are reported as [min - max] proportion of genotyping differences measured for each pair or replicates. Demographic inferences show no filtering on sample quality because more complex extra filtering steps were applied (see main text and supplementary material). Linkage maps also required extra filtering steps described in the main text and supplementary material. The genome scan analysis was based on the exact same set of individuals used in demographic inferences (hence no filter on sample quality). The number of individuals used for linkage maps combines a number of parents (1 or 2) and a number of offspring (55 to 69). Linkage maps involve RAD locus (not individual SNPs). A single SNP per RAD was used in genetic structure analyses and demographic inferences, but not in genome scan analyses where we maximised the number of SNPs present on a RAD locus mapped on our reference map and for which differentiation between species could be calculated in natural populations.

| | Genetic structure | Demographic inference | | Linkage maps | | | | | | | Genome scan | |
|---|---|---|---|---|---|---|---|---|---|---|---|---|
| **Stacks** | | | | | | | | | | | | |
| m | 5 | 3 | | 5 | | | | | | | 5 | |
| M | 2 | 3 | | 5 | | | | | | | 5 | |
| catalog | all samples | all samples | | 7 parents | | | | | | | as linkage maps | |
| n | 5 | 5 | | 5 | | | | | | | 5 | |
| **Filtering** | | | | | | | | | | | | |
| max miss. per SNP | 10% | 30% | | 20% | | | | | | | 50% | |
| max miss. per sample | 15% | - | | 20% | | | | | | | - | |
| **Individuals** | | Britt. | Norm. | Family 1 | | Family 2 | | Family 3 | | Family 4 | Britt. | Norm. |
| *J. albifrons* | 221 | 50 | 40 | 2 + 56 | | 2 + 55 | | | | | 50 | 40 |
| *J. praehirsuta* | 103 | 39 | 38 | | | | | 2 + 69 | | 1 + 59 | 39 | 38 |
| Int. Phenotypes | 25 | | | | | | | | | | | |
| *J. ischiosetosa* | 9 | 9 | | | | | | | | | | |
| **Locus** | | | | Mother | Father | Mother | Father | Mother | Father | Father | | |
| RAD locus | 9718 | 1913 | 1082 | 4712 | 4334 | 4939 | 4639 | 4444 | 4347 | 1512 | | |
| SNPs | 1 per RAD | 1 per RAD | | | | | | | | | 45840 | 35442 |
| SNPs present on our reference map (Family 1 mother) and used in genome scan | | | | | | | | | | | 9028 | 9292 |

### Genetic structure

#### ACP

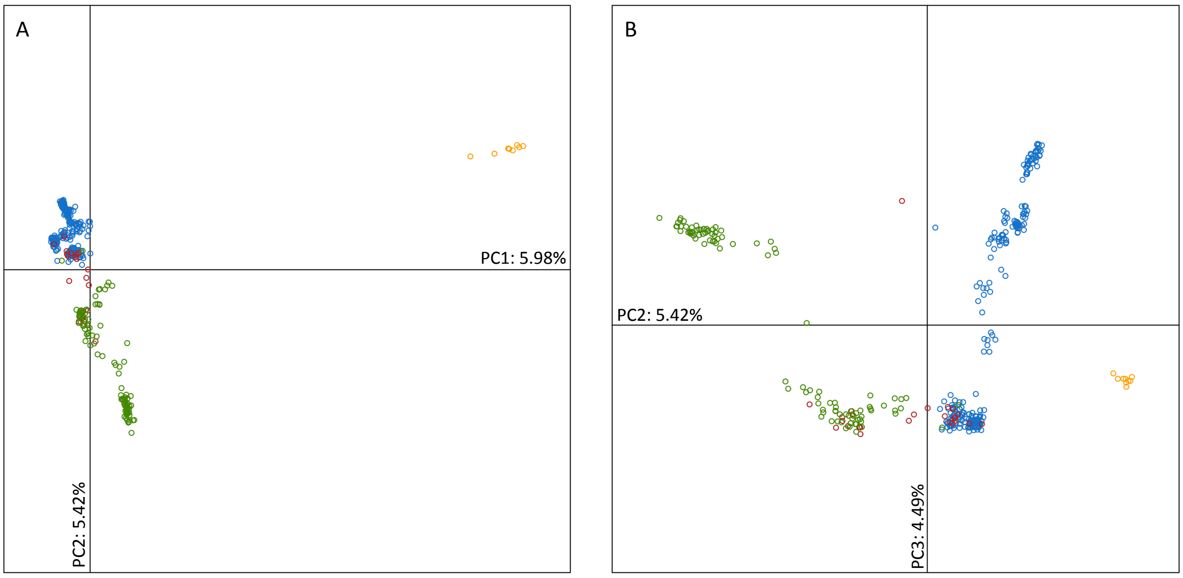


Figure S2 - Principal component analysis based on individual multi-locus genotypes (9718 SNPs derived from ddRAD-seq). The first principal component (PC1, shown in panel A) separates *J. ischiosetosa* individuals (orange) from the two other species (blue: *J. albifrons*, green: *J. praehirsuta*, red: intermediate phenotypes), which are themselves further clustered along PC2. PC3 correlates with the geographic origin of the samples.

#### Treemix

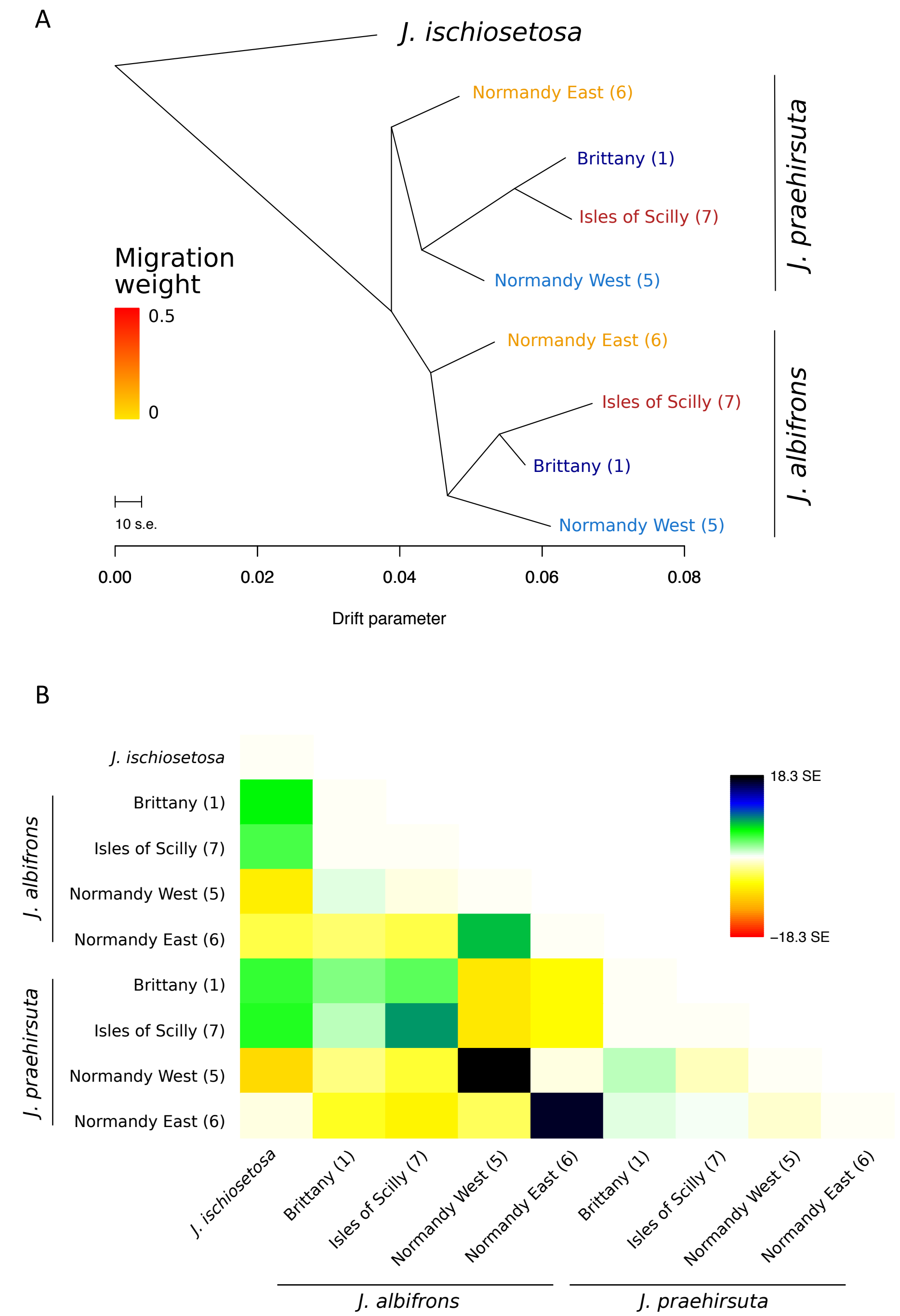


Figure S3 - (A) Maximum-likelihood tree inferred by TreeMix (Pickrell and Pritchard 2012) for four mixed *J. albifrons* / *J. praehirsuta* populations when no migration events are allowed between populations (site numbers and colours as in figure 2). As suggested by the residuals (B), a better fit is obtained when allowing for two migration events between species within sites Normandy West and East (see main text Fig. 2).

### Demographic history of divergence

#### *δaδy*

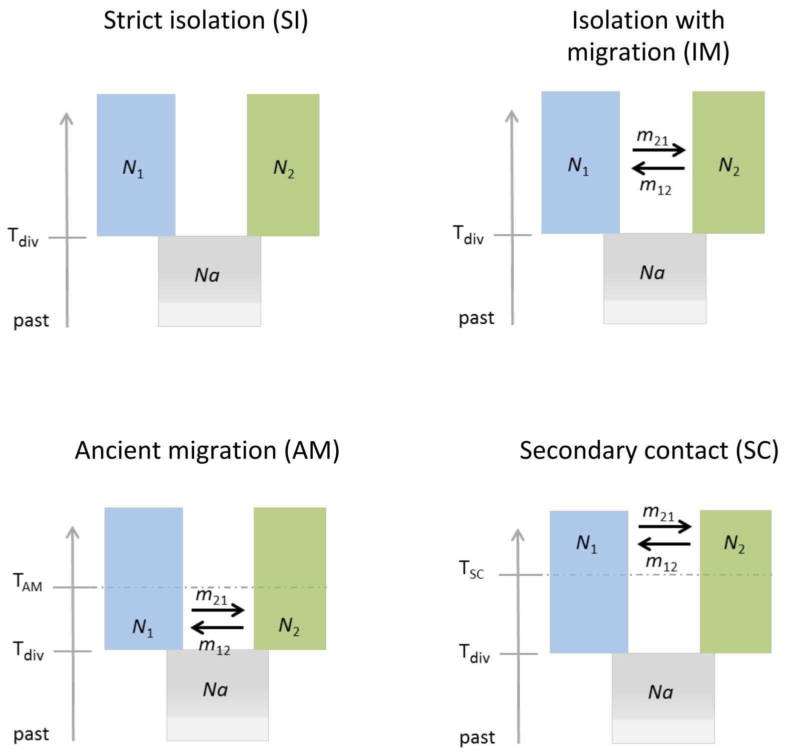


Figure S4 - Demographic models considered for analysing the history of divergence between *J. albifrons* and *J. praehirsuta* in Brittany and Normandy West (main text Fig. 2). The strict isolation model was considered with and without heterogeneous effective population size across loci (SI and SI2N), isolation with migration was considered with and without heterogeneous effective population size and migration rates across the genome (IM, IM2N, and IM2m), and ancient migration and secondary contact models were considered with and without heterogeneous effective population size and migration rates and a combination of these two effects (AM, AM2N, AM2m, AM2N2m, SC, SC2N, SC2m, SC2N2m). Thirteen models were thus considered overall.

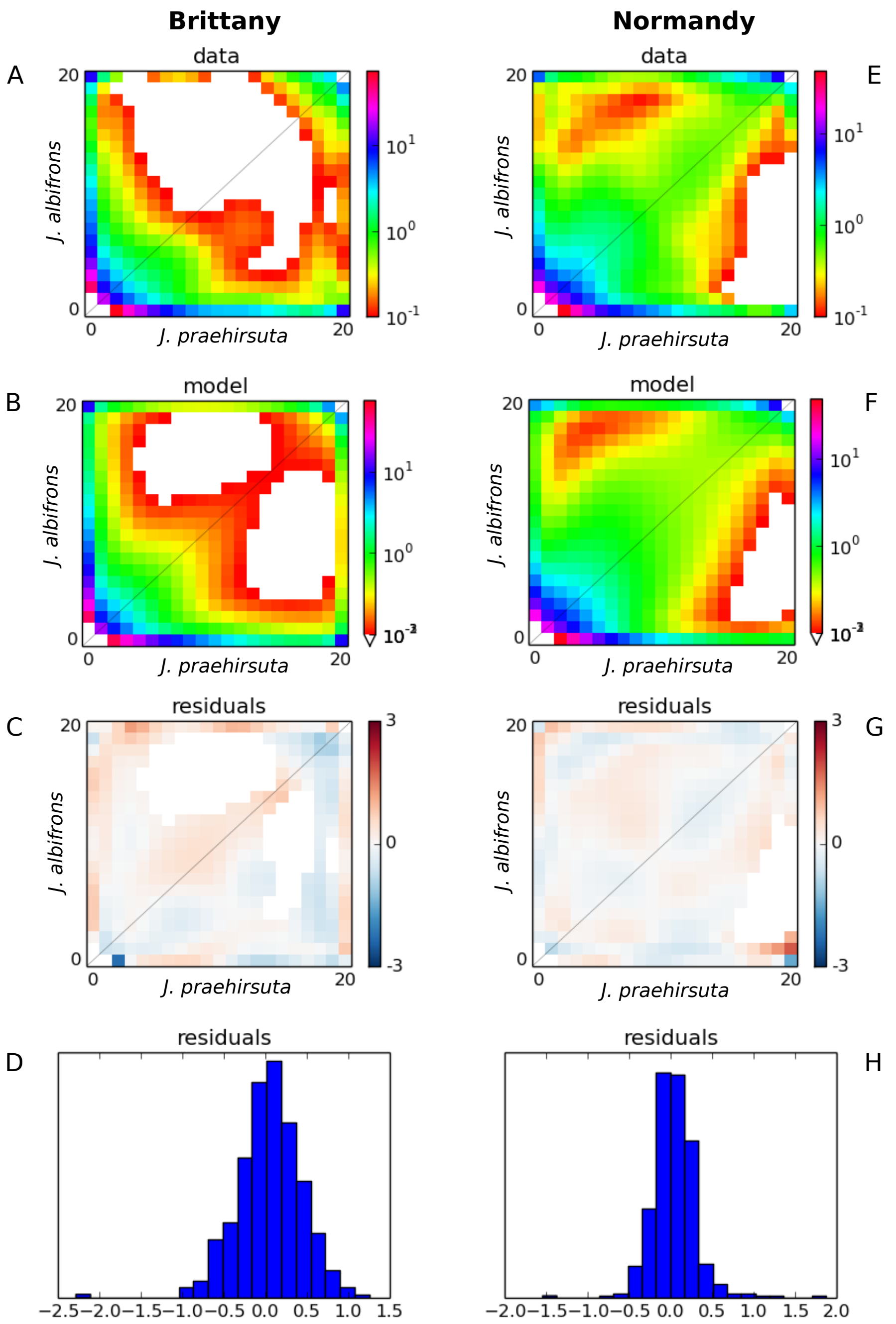


Figure S5 - Comparison between joint allelic frequency spectrums (JAFS) from observed data (A and E), JAFS from the best model (B: SC2N2m and F: SC2m), and the distribution of residuals (C-D and G-H) for region Brittany (A-D) and Normandy West (E-H).

**Table S3:** Estimated values of parameters for the best model (lowest Akaike information criteria AIC, shown in bold) and for all models with a difference in AIC <10 compared to the best model. We also show the values for the SC2m model for region Brittany, for comparison with the results from Normandy West.

| | *AIC* | $\theta$ | $n_1$ | $n_2$ | $Hrf$ | $M_{12}$ | $M_{21}$ | $M'_{12}$ | $M'_{21}$ | $t_S$ | $t_{AM}$ or $t_{SC}$ | $P$ | $Q$ | $O$ |
|---|---|---|---|---|---|---|---|---|---|---|---|---|---|---|
| Brittany | | | | | | | | | | | | | | |
| **SC2N2m** | **569.18** | **16.23** | **5.51** | **6.25** | **0.13** | **2.65** | **1.39** | **0.08** | **0.09** | **4.28** | **0.87** | **0.38** | **0.5** | **0.97** |
| | | | **[4.52;6.50]** | **[4.97;7.52]** | **[0;1]** | **[0;5.83]** | **[0;5.23]** | **[0;0.95]** | **[0;1.08]** | **[3.42;5.15]** | **[0.35;1.39]** | **[0;0.8]** | **[0;1.49]** | **[0.96;0.98]** |
| IM2N | 575.16 | 38.01 | 7.86 | 7.66 | 0.1 | 0.12 | 0.14 | - | - | 4.97 | - | - | 0.48 | 0.98 |
| | | | [6.65;9.07] | [5.42;9.90] | [0;0.68] | [0;1.54] | [0;2.17] | - | - | [3.77;6.18] | - | - | [0;1] | [0.96;0.99] |
| SC2m | 583.84 | 22.96 | 4.54 | 6.64 | | 2.60 | 1.04 | 0.06 | 0.05 | 7.40 | 0.27 | 0.39 | | 0.96 |
| | | | [3.86;5.22] | [6.02;7.27] | | [1.92;3.28] | [0.27;1.81] | [0;0.74] | [0;0.96] | [6.7;8.12] | [0;0.85] | [0.22;0.56] | | [0.95;0.98] |
| Normandy West | | | | | | | | | | | | | | |
| **SC2m** | **684.51** | **103.64** | **0.36** | **1.55** | **-** | **9.93** | **5.81** | **0.39** | **0.54** | **0.78** | **0.094** | **0.67** | **-** | **0.96** |
| | | | **[0;0.95]** | **[1.23;1.88]** | **-** | **[8.57;11.29]** | **[4.89;6.74]** | **[0;1.34]** | **[0;1.14]** | **[0.27;1.30]** | **[0;0.87]** | **[0.39;0.96]** | **-** | **[0.94;0.98]** |
| AM2m | 687.02 | 68.56 | 0.52 | 2.32 | - | 5.12 | 2.17 | 0.18 | 0.13 | 2.63 | 0.00 | 0.72 | - | 0.96 |
| | | | [0;2.18] | [0.87;3.77] | - | [3.11;7.14] | [0.59;3.75] | [0;2.1] | [0;1.61] | [0.31;4.94] | [0.44;0.74] | [0.52;0.92] | - | [0.93;0.99] |
| IM2m | 689.94 | 88.96 | 0.46 | 1.71 | - | 3.96 | 1.95 | 0.00 | 0.13 | 1.6 | - | 0.81 | - | 0.96 |
| | | | [0;1.27] | [0.85;2.58] | - | [3.03;4.90] | [0.59;3.31] | [0;2.38] | [0;1.6] | [0;3.27] | - | [0.69;0.93] | - | [0.94;0.99] |
| SC2N2m | 690.71 | 18.76 | 1.17 | 5.58 | 0.13 | 2.52 | 1.3 | 0.06 | 0.06 | 4.64 | 0.97 | 0.61 | 0.49 | 0.94 |
| | | | [1.09;1.25] | [4;7.16] | [0;0.67] | [1.52;3.52] | [0;4.62] | [0;2.22] | [0;2.13] | [2.6;6.68] | [0.95;0.99] | [0.06;1] | [0;2.24] | [0.9;0.99] |

The table contains for each model: the *AIC* of the model, the estimated genetic diversity $\theta$ of the ancestral population, the scaled effective population size of *J. albifrons* ($n_1$) and *J. praehirsuta* ($n_2$), the Hill-Robertson factor ($Hrf$) corresponding to the reduction in effective population size in some genomic regions, the scaled migration rate from *J. praehirsuta* to *J. albifrons* ($M_{12}$) and in the opposite direction ($M_{21}$), the scaled reduced migration rates corresponding to genomic regions affected by barrier effects ($M'_{12}$ and $M'_{21}$), the time since divergence between the two species ($t_S$), the time since migration has stopped in AM models ($t_{AM}$) or time since migration has resumed after secondary contact ($t_{SC}$), the proportion of the genome that is not affected by barrier effects ($P$), the proportion of the genome that is not affected by the estimated reduction in effective population size ($Q$, where $Hrf$ does not apply), and the proportion of loci for which the ancestral allelic state was properly identified ($O$). All parameters are defined as in (Gutenkunst et al. 2009) and (Rougeux et al. 2017). Bootstrap 95% confidence intervals are given in square brackets.

Limits on the comparison of demographic models across regions:

Comparing absolute values of interspecies migration in the two regions is not straightforward because the best model for Brittany also included an additional factor allowing a fraction of the genome (estimated to 13%, Table S3) to have reduced local effective size due to selection at linked sites. Moreover, the diversity of the ancestral population theta was estimated to be much greater in Normandy than Brittany (Table S3), suggesting either that there were several divergence events between the two species, or that the model cannot completely distinguish the effect of drift from that of gene flow, leading to an overestimation of the ancestral genetic diversity in Normandy.

#### Linkage maps

Linkage maps were built following Stacks manual section 4.4.3 "*De novo* genetic mapping cross", using the seven parents of the families to build the catalog (the mother of family 4 was not genotyped), with Joinmap output format. The genotype data were then filtered in R (based on missing data, Table S2) and passed on to Joinmap (Van Ooijen 2006) to detect segregation distortion. The loci showing highly significant segregation distortion were then filtered out in R. Stacks-populations was then used to generate a vcf data file for each family containing only the filtered set of individuals and RAD loci.

Genetic maps were then constructed using Lep-MAP3 (Rastas 2017). The ParentCall2 module of Lep-MAP3 was used to call parental genotypes, the SeparateChromosomes2 module was used to split the markers into linkage groups, and the JoinSingles2 module added singleton markers when possible. OrderMarkers2 module was then used to order the markers within each linkage group using 30 iterations per group and finally computing genetic distances. The options "informativeMask=13" and "informativeMask=23" were used for male and female maps, respectively, with the modules SeparateChromosomes2 and OrderMarkers2. The phased output data were converted into R/qtl back-cross format corresponding to parental male and female maps.

The maps produced by Lep-MAP3 where then examined, compared, reordered and plotted two-by-two (Tables S4-11) using R functions from the package qtl (Broman et al. 2003) and RCircos (Zhang et al. 2013).

**Table S4**: Summary of the seven linkage maps built for the parents of four F1 families. The map used as a reference for genome scan analyses (Figs. 7, S5, and S6) is that of the first female (*J. albifrons* family 1, highlighted).

| Species | Family | Nb Offspring | Sex | Nb RAD locus | Nb LG |
|---|---|---|---|---|---|
| ***J. albifrons*** | **1** | **56** | **F** | **4712** | **10** |
| | | | M | 4334 | 11 |
| | 2 | 55 | F | 4939 | 11 |
| | | | M | 4639 | 11 |
| *J. praehirsuta* | 3 | 69 | F | 4444 | 12 |
| | | | M | 4347 | 11 |
| | 4 | 59 | M | 1512 | 12 |

**Table S5**: Summary statistics of the linkage map built for the mother of family 1 (*J. albifrons* female)

| LG | n.mar | length | ave.spacing | max.spacing |
|---|---|---|---|---|
| 1 | 637 | 71.65 | 0.11 | 9.03 |
| 2 | 552 | 71.49 | 0.13 | 5.38 |
| 3 | 267 | 46.5 | 0.17 | 7.19 |
| 4 | 220 | 46.74 | 0.21 | 12.77 |
| 5a-b | 571 | 103.73 | 0.18 | 7.19 |
| 5c | 367 | 55.39 | 0.15 | 3.58 |
| 6 | 574 | 78.65 | 0.14 | 5.38 |
| 7 | 592 | 62.55 | 0.11 | 5.38 |
| 8 | 483 | 55.42 | 0.11 | 5.38 |
| 9 | 449 | 67.92 | 0.15 | 5.38 |
| overall | 4712 | 660.05 | 0.14 | 12.77 |

**Table S6**: Summary statistics of the linkage map built for the father of family 1 (*J. albifrons* male)

| LG | n.mar | length | ave.spacing | max.spacing |
|---|---|---|---|---|
| 1 | 407 | 62.55 | 0.15 | 5.38 |
| 2 | 649 | 57.29 | 0.09 | 9.02 |
| 3 | 242 | 58.98 | 0.24 | 3.58 |
| 4 | 209 | 55.60 | 0.27 | 10.88 |
| 5a | 415 | 53.66 | 0.13 | 7.19 |
| 5b | 145 | 46.62 | 0.32 | 10.88 |
| 5c | 349 | 48.28 | 0.14 | 5.38 |
| 6 | 615 | 73.31 | 0.12 | 5.38 |
| 7 | 559 | 76.88 | 0.14 | 7.19 |
| 8 | 337 | 10.72 | 0.03 | 1.79 |
| 9 | 407 | 67.94 | 0.17 | 5.38 |
| overall | 4334 | 611.84 | 0.14 | 10.88 |

**Table S7**: Summary statistics of the linkage map built for the mother of family 2 (*J. albifrons* female)

| LG | n.mar | length | ave.spacing | max.spacing |
|---|---|---|---|---|
| 1 | 655 | 78.49 | 0.12 | 11.09 |
| 2 | 638 | 69.41 | 0.11 | 11.09 |
| 3 | 278 | 42.04 | 0.15 | 9.19 |
| 4 | 216 | 38.2 | 0.18 | 3.64 |
| 5a | 399 | 47.41 | 0.12 | 9.19 |
| 5b | 145 | 45.57 | 0.32 | 9.19 |
| 5c | 384 | 58.24 | 0.15 | 5.48 |
| 6 | 571 | 89.17 | 0.16 | 3.64 |
| 7 | 591 | 76.45 | 0.13 | 5.48 |
| 8 | 650 | 65.56 | 0.1 | 5.48 |
| 9 | 412 | 80.11 | 0.19 | 5.48 |
| overall | 4939 | 690.65 | 0.14 | 11.09 |

**Table S8**: Summary statistics of the linkage map built for the father of family 2 (*J. albifrons* male)

| LG | n.mar | length | ave.spacing | max.spacing |
|---|---|---|---|---|
| 1 | 387 | 56.42 | 0.15 | 5.48 |
| 2 | 559 | 52.77 | 0.09 | 5.48 |
| 3 | 311 | 52.79 | 0.17 | 5.48 |
| 4 | 211 | 52.78 | 0.25 | 5.48 |
| 5a | 453 | 61.88 | 0.14 | 5.48 |
| 5b | 165 | 38.21 | 0.23 | 3.64 |
| 5c | 377 | 38.21 | 0.1 | 3.64 |
| 6 | 589 | 83.71 | 0.14 | 5.48 |
| 7 | 598 | 81.89 | 0.14 | 3.64 |
| 8 | 525 | 72.79 | 0.14 | 3.64 |
| 9 | 464 | 72.84 | 0.16 | 7.32 |
| overall | 4639 | 664.28 | 0.14 | 7.32 |

**Table S9:** Summary statistics of the linkage map built for the mother of family 3 (*J. praehirsuta* female)

| LG | n.mar | length | ave.spacing | max.spacing |
|---|---|---|---|---|
| 1 | 629 | 42.13 | 0.07 | 7.3 |
| 2 | 503 | 72.57 | 0.14 | 7.3 |
| 3 | 180 | 34.81 | 0.19 | 2.9 |
| 4 | 214 | 46.48 | 0.22 | 8.79 |
| 5a | 348 | 56.6 | 0.16 | 5.82 |
| 5b | 176 | 24.65 | 0.14 | 2.9 |
| 5c | 427 | 33.36 | 0.08 | 4.36 |
| 6a | 363 | 44.95 | 0.12 | 4.36 |
| 6b | 202 | 23.2 | 0.12 | 2.9 |
| 7 | 578 | 71.12 | 0.12 | 7.3 |
| 8 | 356 | 58.03 | 0.16 | 5.82 |
| 9 | 468 | 75.41 | 0.16 | 4.36 |
| overall | 4444 | 583.31 | 0.13 | 8.79 |

**Table S10:** Summary statistics of the linkage map built for the father of family 3 (*J. praehirsuta* male)

| LG | n.mar | length | ave.spacing | max.spacing |
|---|---|---|---|---|
| 1 | 347 | 72.99 | 0.21 | 13.35 |
| 2 | 499 | 66.73 | 0.13 | 4.36 |
| 3 | 266 | 50.81 | 0.19 | 7.3 |
| 4 | 202 | 58.13 | 0.29 | 8.79 |
| 5a | 354 | 59.45 | 0.17 | 4.36 |
| 5b-c | 463 | 66.86 | 0.14 | 10.29 |
| 6a | 369 | 62.46 | 0.17 | 8.78 |
| 6b | 239 | 53.69 | 0.23 | 4.36 |
| 7 | 613 | 82.67 | 0.14 | 4.36 |
| 8 | 417 | 34.84 | 0.08 | 5.82 |
| 9 | 578 | 66.76 | 0.12 | 5.82 |
| overall | 4347 | 675.4 | 0.16 | 13.35 |

**Table S11**: Summary statistics of the linkage map built for the father of family 4 (*J. praehirsuta* male)

| LG | n.mar | length | ave.spacing | max.spacing |
|---|---|---|---|---|
| 1 | 148 | 85.6 | 0.58 | 12.09 |
| 2 | 191 | 52.75 | 0.28 | 8.56 |
| 3 | 74 | 33.93 | 0.46 | 3.4 |
| 4 | 98 | 65.48 | 0.68 | 19.59 |
| 5a | 112 | 37.88 | 0.34 | 15.76 |
| 5b-c | 133 | 44.21 | 0.33 | 8.56 |
| 6a | 154 | 46.04 | 0.3 | 10.31 |
| 6b | 64 | 52.78 | 0.84 | 8.56 |
| 7a | 84 | 49.6 | 0.6 | 13.91 |
| 7b | 73 | 33.92 | 0.47 | 3.4 |
| 8 | 136 | 88.31 | 0.65 | 17.65 |
| 9 | 245 | 85.11 | 0.35 | 12.1 |
| overall | 1512 | 675.61 | 0.45 | 19.59 |

#### Identification of sex chromosomes

To identify sex chromosomes (and not identify each and every locus sitting on sex chromosomes), we used only the most informative markers, that is, the RAD-locus that are heterozygous in the mother of a given family and homozygous in the father. By convention, these locus are coded as lm x ll, meaning that the mother of a cross has alleles l and m, and the father is homozygous for allele l (although note that in our Joinmap-formatted .loc datasets, because the first parent identity was that of the father followed by that of the mother, the lm x ll genotypes were in fact coded as nn x np, where the mother is np).

With this type of markers, if the locus is autosomal, then genotype frequencies do not differ in the male and female offspring (on average half of the individuals will be ll, half lm, regardless of their sex). By contrast, if a locus is located on a non-recombining region of ZW sex chromosomes, then the distribution of genotypes will be drastically different. If allele m is located on the Z chromosome of the female, then all female offspring will be homozygous ll and all male offspring will be heterozygous lm. Similarly, if allele m is located on the W of the mother, then all female offspring will have the lm genotype and all male offspring will have ll genotypes. Hence, all locus that follow a segregation pattern where all offspring of one sex are homozygous ll and all offspring of the other sex are heterozygous lm are compatible with and only with ZW segregation.

We identified all locus following this pattern in each of our three families (excluding the *J. praehirsuta* family for which we had only the father genotypes, see main text). We found 275 to 335 markers following this pattern (table Sx), and they all appeared to be located on a unique, common linkage group across all our 3 mothers (they cannot be present on the father's map because these loci are homozygous in the father), unequivocally pointing towards a unique pair of sex chromosomes (here noted LG1). Moreover, all sex-linked RAD locus identified were found to collocate nearly all at a unique position within each map (Table Sx), meaning that they were all included in a non-recombining region of unknown physical size that contains the sex-determining gene.

While we could not formally identify the sex chromosomes using the recombination map built for the *J. praehirsuta* male of family 4 (because this map was built using only snps that were heterozygous in the male and homozygous in the female parent), there is no reason to think that its linkage group corresponding to LG1 in the maps obtained for the six other individuals is not also corresponding to the sex chromosomes.

**Table S12**: Identification of sex chromosomes using informative locus from the linkage maps. For each family: number of locus of type lm x ll, number of such locus that are strictly heterozygous in all offspring of one sex and homozygous in the other sex, and position of these sex-linked locus (linkage group and position within this LG)

| Family | lm x ll | sex-linked | linkage group | position on LG |
|---|---|---|---|---|
| *J. albifrons* family 1 | 2535 | 307 | LG1 | 44.749 [a] |
| *J. albifrons* family 2 | 2595 | 335 | LG1 | 43.738 |
| *J. praehirsuta* family 3 | 2459 | 275 | LG1 | 39.232 [b] |

[a] all locus were localized at the same map position, except one locus at position 46.736
[b] all locus were localized at the same map position, except one locus at position 40.682

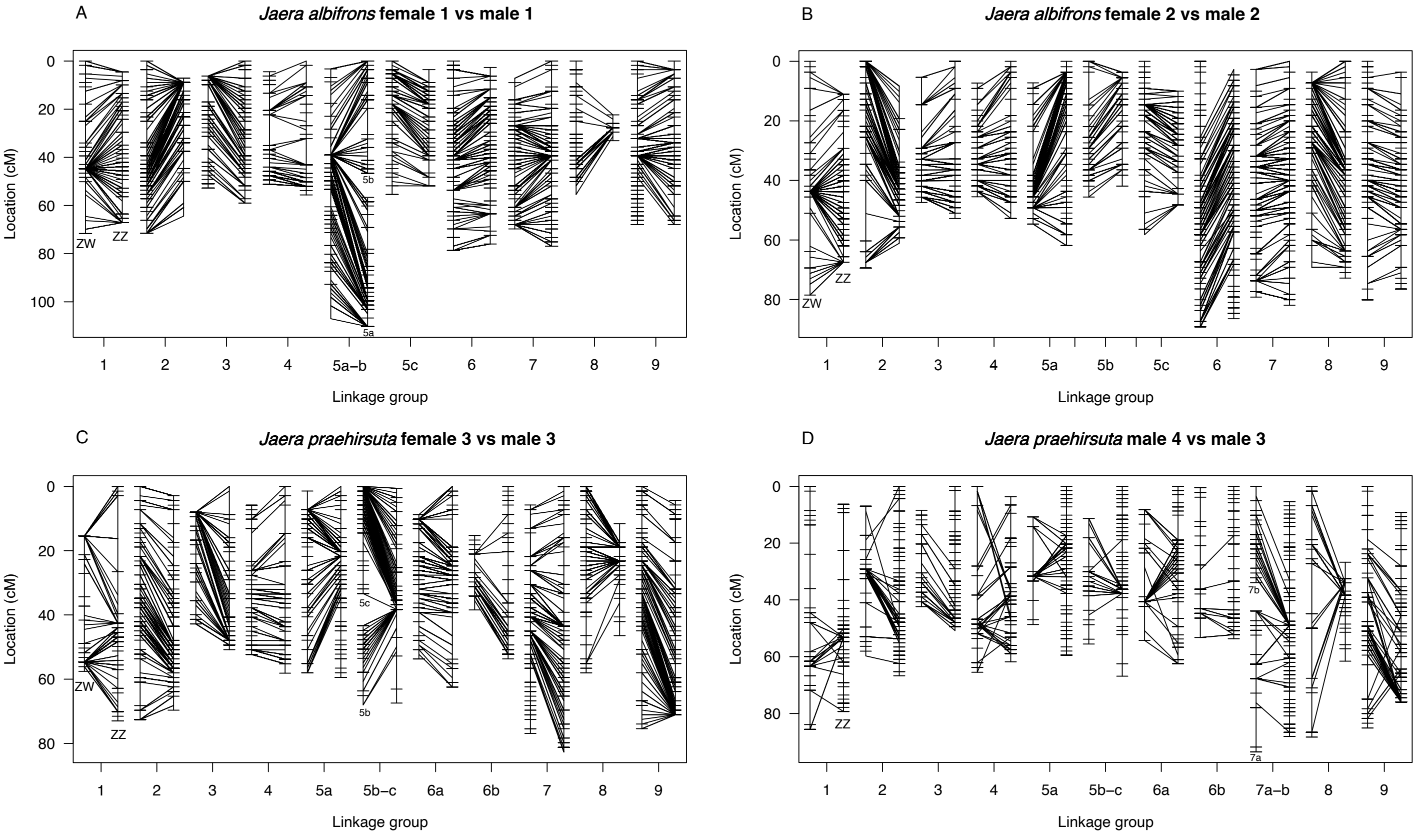


Figure S6 - Intraspecific dual comparisons of the linkage maps obtained for seven individuals (see Table S4). Panels A and B compare linkage maps for four *J. albifrons* (maternal and paternal maps on the left and right within each panel, respectively). Panel C shows the maps obtained for *J. praehirsuta* family 3, and panel D shows the male map of family 4 (left) against the map obtained for family 3 (right, already shown in panel C right). Within each panel, the RAD locus that were found in common between the two maps compared are connected by a straight line. These plots can only show the markers that were found to be on the same linkage groups across two maps, except for the linkage groups 5a, b, c, and 7a,b, which correspond to intra-specific Robertsonian rearrangements (in these cases, when two linkage groups are shown simultaneously, they are arbitrarily separated here by a 10 cM gap for visualization). Additional chromosomal rearrangements between species are shown in figures 6 and 7 and in supplementary figure S6. The sex chromosomes independently identified for the seven adults all corresponded to LG1.

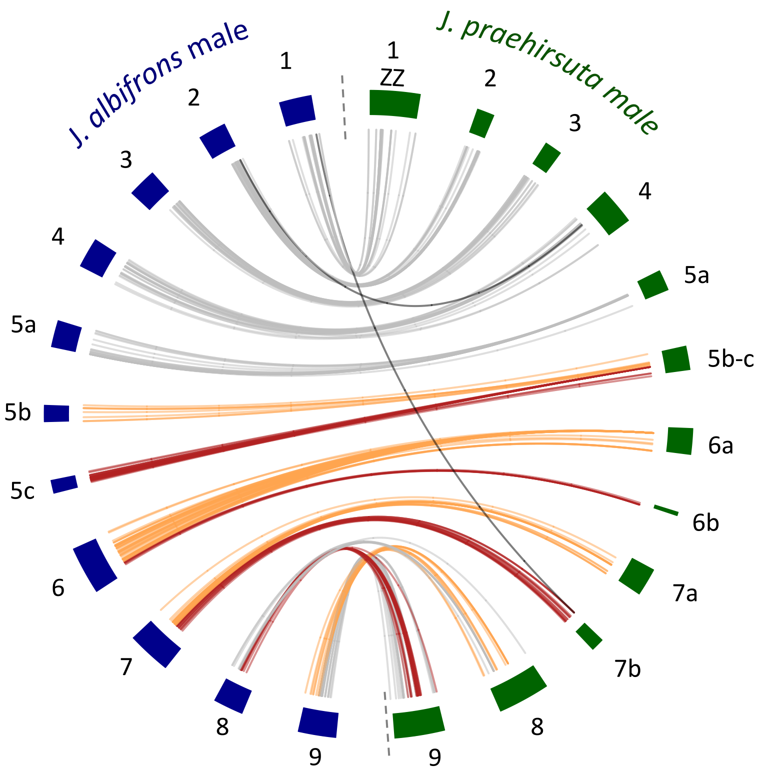


Figure S7 - Genome synteny and collinearity based on linkage maps from male *Jaera albifrons* (family 2) and male *J. praehirsuta* (family 4). Numbered rectangles correspond to linkage groups (LGs). The LGs corresponding to the sex chromosomes were given the number 1, but sex-linkage could not be formally identified in the case of this *J. praehirsuta* male (for lack of genetic data for the mother of family 4). As in main text figure 6, each SNP common to two maps is shown in grey when it occurs on the same LG in both maps, and in orange or red when it occurs on different LGs. Putative mapping errors are shown in black. This figure confirms the rearrangements between species observed when using families 1-3 (which did not lack the mother's genotypes and were thus based on a much larger number of snps, see main text): fusion/scissions of LG5 and LG6, and reciprocal translocation of LGs8-9. In addition, it suggests another rearrangement (fusion/scission) within *J. praehirsuta* involving LG7 (here divided into two groups in the recombination map obtained for the *J. praehirsuta* male).

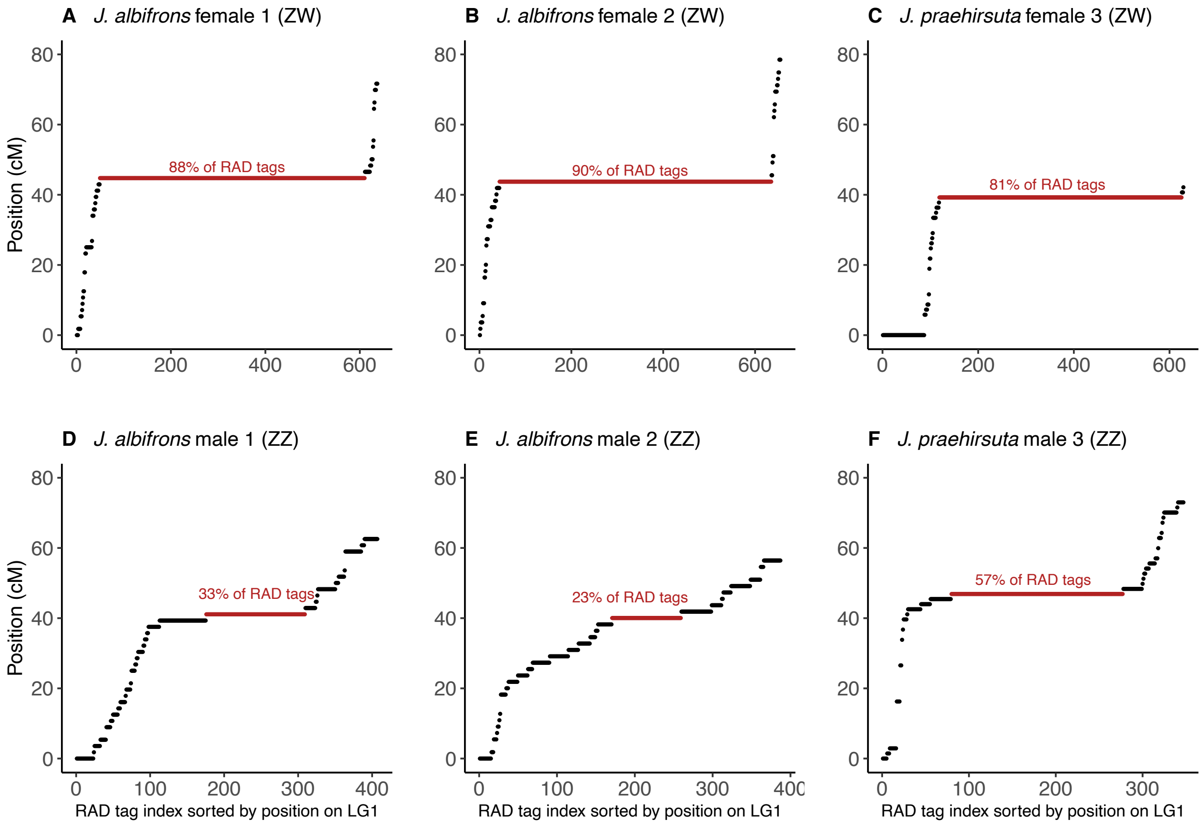


Figure S8 - Distribution of RAD sequences on linkage group 1 (sex chromosomes) in the three families for which a map was obtained for the mother and the father (families 1-3 in table S4). Here, RAD sequences are sorted according to their position on the linkage group (in cM). A region of low or null recombination can be identified from the concentration of markers (shown in red). Panels A-C show the variation in marker density in females. Assuming that RAD tags are essentially randomly distributed, we see that ZW recombination is absent in a large region (80 to 90% of the sex chromosomes). Interestingly, a similar, although less clear-cut pattern is also visible in males (panels D-F), meaning that ZZ recombination seems also reduced in a central region of these chromosomes. The red zone is defined here as a portion of the linkage map aggregating the largest number of RAD-tags at a unique position.

### Genome scans

#### Simulations

We used ms (Hudson 2002) to simulate bi-allelic loci evolving neutrally under a secondary contact scenario. Results from *δaδi* showed that a scenario of secondary contact with heterogeneous migration and heterogeneous effective population size along the genome (SC2N2m) was the best fit to predict the allelic frequency spectrum of *Jaera albifrons* and *J. praehirsuta* in region Brittany (Fig. S4). However, we chose to simulate a simplified model including only heterogeneous migration (SC2m, also a good fit, Table S3) so that the results of the simulations are directly comparable to that of region Normandy (for which SC2m was also the model most closely mimicking the observations). Based upon parameter values for the SC2m in Brittany and Normandy (Table S3), we used the following code for simulating data in Brittany and Normandy, respectively:

#ms 178 100000 -t 22.963 -s 1 -I 2 100 78 0 -ma X 5.201 2.078 X -n 1 4.543 -n 2 6.644 -eM 0.133 0 -ej 3.702 2 1 -eN 3.702 1 >> br_100000_m.txt

#ms 158 100000 -t 103.642 -s 1 -I 2 80 78 0 -ma X 19.86 11.622 X -n 1 0.363 -n 2 1.551 -eM 0.047 0 -ej 0.392 2 1 -eN 0.392 1 >> no_100000_m.txt

The parameterization takes into account the difference in scaling factors between *δaδi* (time and migration scaled by 2Nref) and ms (time and migration scaled by 4Nref). These simulations were run twice to produce 200 000 SNPs, which were then converted in R for estimating diversity across ($H_T$) and differentiation between species ($F_{ST}$) at each locus (Fig. S7).

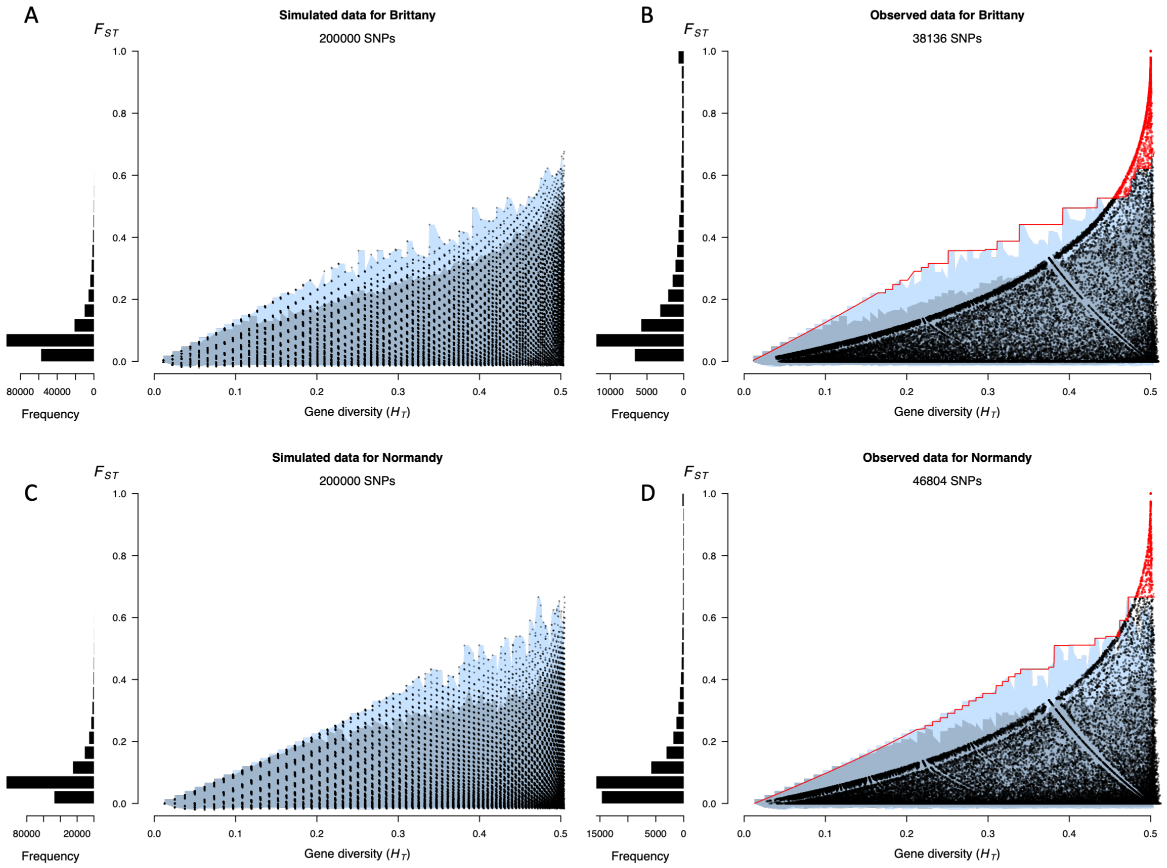


Figure S9 - Panels A and C: genetic differentiation ($F_{ST}$) between *J. albifrons* and *J. praehirsuta* at 200000 simulated SNPs using a neutral model parameterized following the demographic parameters inferred from δaδi in Brittany (A) and Normandie (B) as a function of total gene diversity ($H_T$). The two shades of blue show the envelopes containing 95 % and 100 % of the highest $F_{ST}$ values (simulated data), respectively. Histograms on the left present the distribution of locus-by-locus $F_{ST}$ values. Panels B and D: observed genetic differentiation ($F_{ST}$) between *J. albifrons* and *J. praehirsuta* in Brittany (B, 38136 SNPs) and Normandy (D, 46804 SNPs). The blue envelopes are inherited from the simulations shown in left panels. Red dots show the $F_{ST}$ higher than any value obtained through simulations (the corresponding SNPs are shown in red in main text Fig. 4).

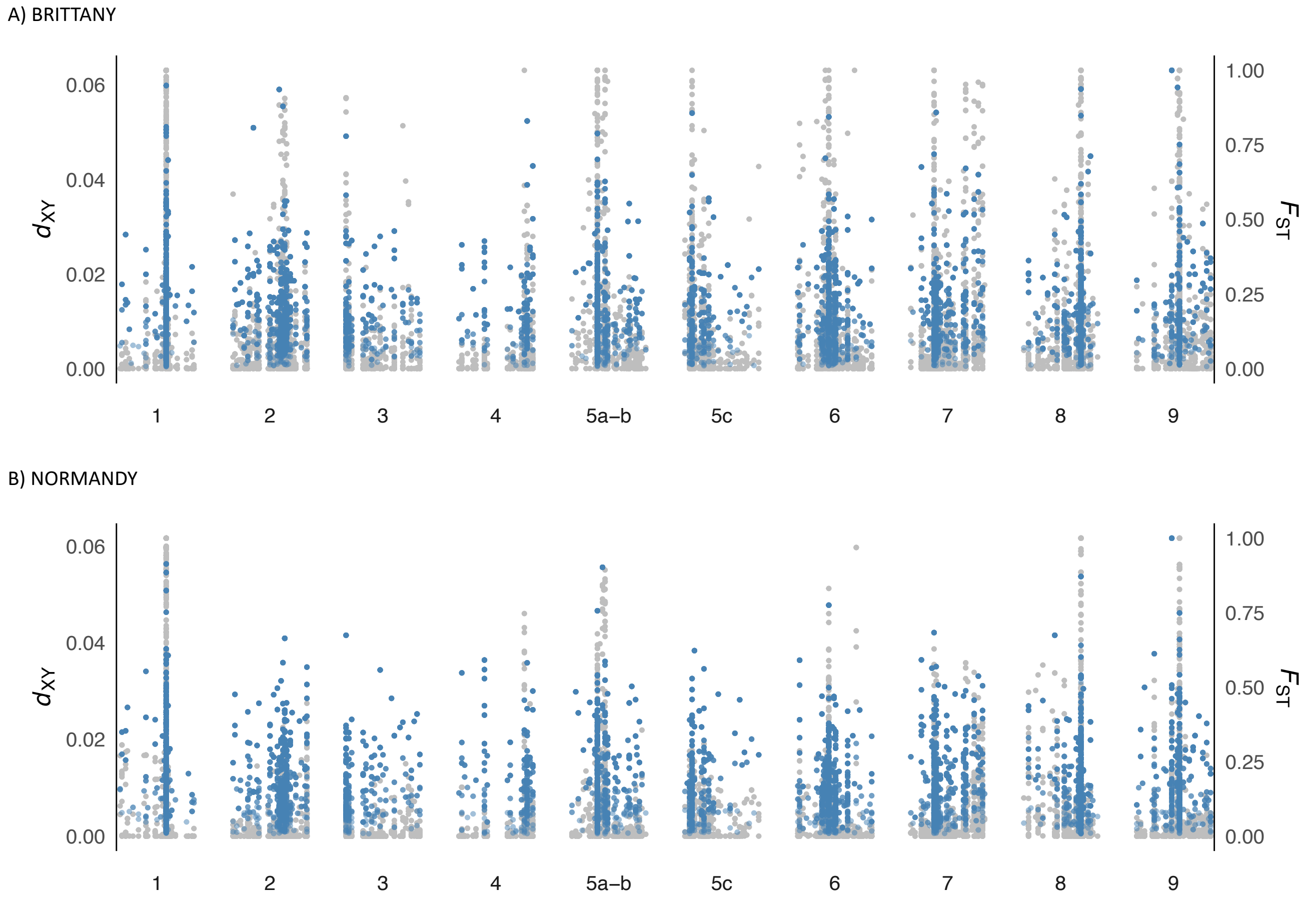


Figure S10 - Absolute divergence $d_{XY}$ (in blue, primary Y-axis), between *J. albifrons* and *J. praehirsuta* in region Brittany (Fig. 2 site 1, 2925 RAD-tag sequences) and region Normandy West (Fig. 2 site 5, 3288 RAD-tags). The grey points in the background give the $F_{ST}$ values (secondary Y-axis) shown in main text figure 9.

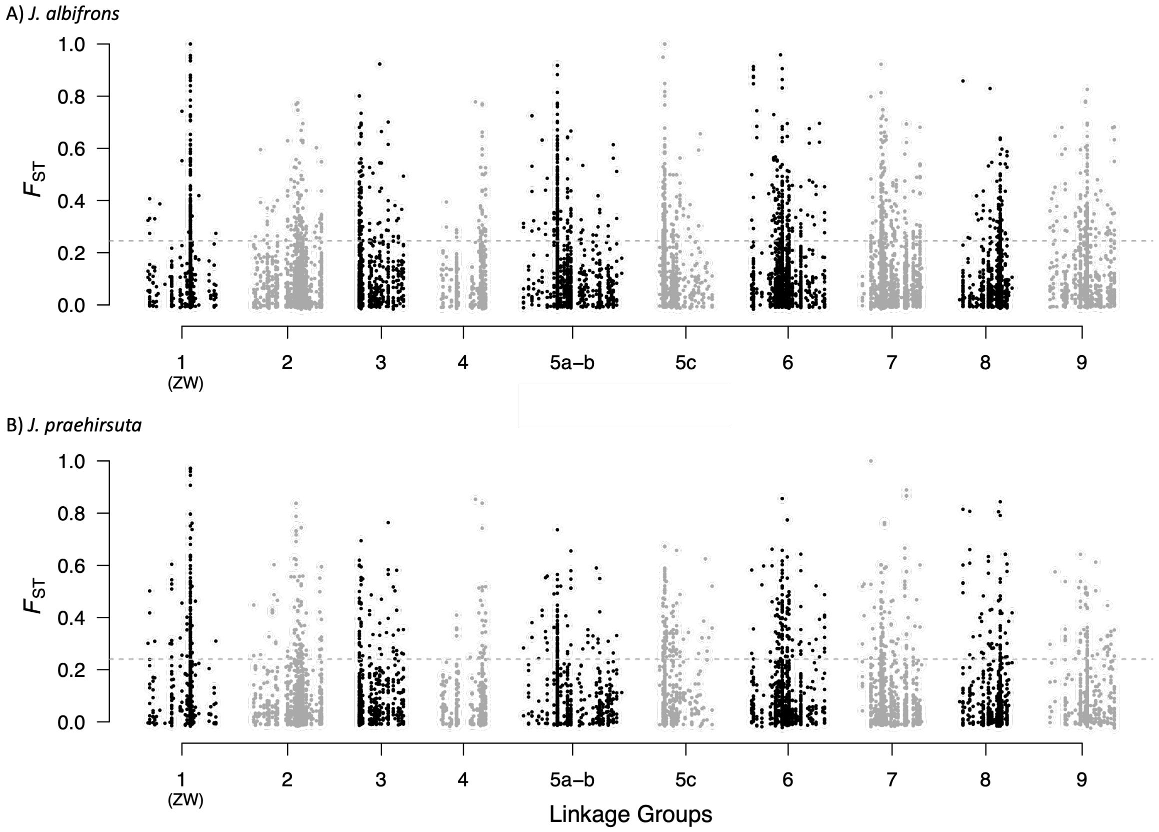


Figure S11 - Distribution of genetic differentiation ($F_{ST}$) between geographic regions (Brittany vs Normandy) in either *J. albifrons* (A) or *J. praehirsuta* (B). Each point represents the $F_{ST}$ value between species measured at one SNP (here 11152 and 7667 SNPs, respectively). The reference map used here is that of a female *J. albifrons* (maternal map of family 1, shown in light blue in main text Fig. 6A). The horizontal dashed lines give the genome-wide $F_{ST}$ (i.e. calculated over all SNPs) in each species.

### QTL analysis

#### Families

To identify and map the loci potentially linked to the genes encoding male secondary sexual traits, we used an independent dataset obtained from a collection of 15 backcross families designed to show high phenotypic variation in male courtship traits (Table S13).

**Table S13**: Backcross families used for QTL analysis. One of the parents of each family is an F1 hybrid previously obtained in the laboratory from a controlled cross between *J. albifrons* and *J. praehirsuta* individuals. The number of offspring corresponds to the number of male progeny reared until their secondary sexual characteristics could be described.

| Family | Mother | Father | Nb offspring |
|---|---|---|---|
| G1-1 | F1 | *J. praehirsuta* | 8 |
| G1-2 | F1 | *J. praehirsuta* | 14 |
| G2-1 | F1 | *J. albifrons* | 14 |
| G2-2 | F1 | *J. albifrons* | 7 |
| G3 | F1 | *J. praehirsuta* | 11 |
| G4-1 | F1 | *J. praehirsuta* | 15 |
| G4-2 | F1 | *J. praehirsuta* | 22 |
| G5 | F1 | *J. praehirsuta* | 18 |
| G6-1 | F1 | *J. albifrons* | 6 |
| G6-2 | F1 | *J. albifrons* | 3 |
| G7-1 | F1 | *J. albifrons* | 4 |
| G7-2 | F1 | *J. albifrons* | 7 |
| G8 | F1 | *J. albifrons* | 6 |
| G9 | *J. praehirsuta* | F1 | 10 |
| G10 | *J. praehirsuta* | F1 | 5 |

#### Phenotypic traits

Overall, 150 male offspring were reared until they secondary sexual characters could be analysed (rearing conditions described in Ribardière et al. 2021). These traits included the presence/absence of a carpal lobe on peraeopods P6 and P7, the number of sexual setae on P1-P4 and P6-P7, and the number of spines on P6-P7 (main text Fig. 1). The length of each individual was also measured as in Ribardière et al. (2021).

Because the number of setae and spines correlate with the total length of individuals (Prunus 1968), these values were regressed against individual length, and the standardized residuals of these regression models were used in QTL analyses. Numbers of sexual setae had zero-inflated distributions, because a majority of males tended to resemble *J. albifrons* males (no sexual setae on P1-4) or *J. praehirsuta* males (no sexual setae on P6-P7). Since these zero-inflated distributions could not be satisfactorily corrected, for P1-4 and P6-7, we considered the presence/absence of sexual setae and then the number of setae when present.

**Table S14**: Male secondary sexual traits used in courtship. The number of setae on P1-4 and P6-7 had zero-inflated distributions due to the contrasting phenotypes between species. These traits were thus treated first as a binomial variable (presence-absence) and second as a quantitative variable when present (number of setae > 0).

| Trait | Description | Type | Range / values | Size regression method for continuous variables |
|---|---|---|---|---|
| P1 setae presence | Presence of curved setae on P1 | Binomial | Presence - absence | |
| Nb of P1 setae | Number of curved setae on P1, if >0 | Integer | 1 - 51 | Normalized residuals of linear regression |
| P2 setae presence | Presence of curved setae on P2 | Binomial | Presence - absence | |
| Nb of P2 setae | Number of curved setae on P2, if >0 | Integer | 1 - 45 | Normalized residuals of linear regression |
| P3 setae presence | Presence of curved setae on P3 | Binomial | Presence - absence | |
| Nb of P3 setae | Number of curved setae on P3, if >0 | Integer | 1 - 51 | Normalized residuals of linear regression |
| P4 setae presence | Presence of curved setae on P4 | Binomial | Presence - absence | |
| Nb of P4 setae | Number of curved setae on P4, if >0 | Integer | 1 - 37 | Normalized residuals of linear regression |
| P6 setae presence | Presence of curved setae on P6 | Binomial | Presence - absence | |
| Nb of P6 setae | Number of setae on P6, if > 0 | Integer | 1 - 30 | Normalized residuals of linear regression |
| P6 lobe presence | Presence of a carpal lobe on P6 | Binomial | Presence - absence | |
| Nb of P6 spines | Number of spines on P6 | Integer | 0 - 3 | Normalized residuals of Poisson family GLM |
| P7 setae presence | Presence of setae on P7, if > 0 | Binomial | Presence - absence | |
| Nb of P7 setae | Number of setae on P7 carpus, if > 0 | Integer | 1 - 20 | Normalized residuals of linear regression |
| P7 lobe presence | Presence of a carpal lobe on P7 | Binomial | Presence - absence | |
| Nb of P7 spines | Number of spines on P7 | Integer | 0 - 4 | Normalized residuals of Poisson family GLM |

The final set of phenotypic traits used in QTL analysis (Table S14) thus included 8 binomial variables (presence/absence of setae on P1-4 and P6-7, and presence/absence of a carpal lobe on P6-7) and the normalized residuals of 8 continuous variables (number of setae when >0 on P1-4 and P6-7, and number of spines on P6-7). As shown in Table S14, the number of spines on P6 and P7 was best regressed against total length by using a GLM with a Poisson distribution, while a linear model was adequate for the remaining six continuous variables (number of setae when present).

#### Genetic map

An average, consensus map was built using the 15 backcross families with Lep-MAP3 as described above in section "Linkage maps" except that the option “sexAveraged=1” was used with the module OrderMarkers2. The phased output data were converted into F2 type format with three genotypes AA, AB and BB used for QTL mapping.

QTLs were identified using the R packages qtl (Broman et al. 2003) and qtl2 (Broman et al. 2019). The scanone function, with the Haley-Knott regression method, was used to perform interval mapping in R/qtl. A genome scan using a linear mixed model (LMM), accounting for relationships among individuals using a random polygenic effect, was also performed using the function scan1. LOD (Logarithm of the odds) significances were ensured with 1000 permutation tests. The percentage of variance explained by a QTL was assessed with analysis of variance using type III sums of squares using the fitqtl function. Confidence intervals were calculated as Bayesian credible intervals using bayesesint function with a probability of coverage of 0.95. Finally, marker regression was implemented in the function scanone with method =”mr”.

#### Location of QTL on reference linkage map

The interval mapping method yielded several genomic regions that were significantly associated with male sexual traits. To locate these regions on our reference map (and then with genetic differentiation between species, as shown in main text Fig. 8), we identified the RAD loci that were located exactly at these significant genomic regions in the QTL analysis and then searched for these loci on our reference map (female *J. albifrons*, family 1). If such a RAD locus was not present on the reference map, we used its position on the six other linkage maps (Table S5-S11 and Fig. S5) to identify linked RADs (at the exact same position) that were also present on the reference map.

**Table S15**: Results from the interval mapping QTL analysis. Here the linkage groups (LG) are named accordingly to our reference linkage map (*J. albifrons* family 1 female, Table S5). The confidence interval limits for the position of each QTL are given in columns CI lower limit and CI upper limit. The QTL positions reported in main text Fig. 8 were deduced from the position of RAD loci found in common between these QTL analysis results and the reference linkage map used in all other analyses.

| Trait | Position on QTL linkage map | | | | | | Corresponding position on reference map | |
|---|---|---|---|---|---|---|---|---|
| | LG | pos | lod | CI lower limit | CI upper limit | LOD thresh 5% | LG | pos |
| P1 setae presence | 11 | 38.92 | 9.40 | 18.90 | 41.30 | 3.50 | 1 | 37.6 to 48.3 |
| P2 setae presence | 11 | 38.92 | 7.68 | 18.90 | 41.30 | 3.53 | 1 | 37.6 to 48.3 |
| Nb of P2 setae | 11 | 25.80 | 4.04 | 20.65 | 30.57 | 3.49 | 1 | 44.7 |
| P3 setae presence | 11 | 38.92 | 10.08 | 18.90 | 41.30 | 3.57 | 1 | 37.6 to 48.3 |
| Nb P3 setae | 11 | 25.80 | 3.67 | 20.65 | 30.57 | 3.51 | 1 | 44.7 |
| P4 setae presence | 11 | 39.95 | 4.73 | 0 | 47.63 | 3.54 | 1 | 44.7 |
| P6 setae presence | 11 | 39.27 | 4.82 | 18.90 | 47.63 | 3.56 | 1 | 37.6 to 48.3 |
| P6 lobe presence | 11 | 40.30 | 5.68 | 18.90 | 41.97 | 3.51 | 1 | 44.7 |
| P7 setae presence | 11 | 39.27 | 5.23 | 8.96 | 47.30 | 3.48 | 1 | 37.6 to 48.3 |
| Nb of P7 spines | 11 | 25.80 | 8.61 | 25.80 | 30.57 | 3.52 | 1 | 44.7 |
| P7 lobe presence | 11 | 40.30 | 7.04 | 18.90 | 44.63 | 3.54 | 1 | 44.7 |
| P1 setae presence | 10 | 31.62 | 4.33 | 28.62 | 32.95 | 3.50 | 4 | 43.2 |
| Nb of P1 setae | 10 | 7.53 | 4.09 | 1.26 | 32.62 | 3.51 | 4 | 10.7 |
| Nb P2 setae | 10 | 20.20 | 3.73 | 3.17 | 89.41 | 3.49 | 4 | 44.9 |
| Nb of P7 spines | 6 | 25.79 | 4.53 | 18.50 | 32.49 | 3.52 | 5a-b | 37.5 to 50.1 |

Broman, K. W., D. M. Gatti, P. Simecek, N. A. Furlotte, P. Prins, Ś. Sen, B. S. Yandell, and G. A.

Churchill. 2019. R/qtl2: Software for Mapping Quantitative Trait Loci with High-Dimensional Data and Multiparent Populations. Genetics 211:495–502.

Broman, K. W., H. Wu, S. Sen, and G. A. Churchill. 2003. R/qtl: QTL mapping in experimental crosses. Bioinforma. Oxf. Engl. 19:889–890.

Catchen, J., P. A. Hohenlohe, S. Bassham, A. Amores, and W. A. Cresko. 2013. Stacks: an analysis tool set for population genomics. Mol. Ecol. 22:3124–3140.

Catchen, J. M., A. Amores, P. Hohenlohe, W. Cresko, and J. H. Postlethwait. 2011. Stacks: Building and Genotyping Loci De Novo From Short-Read Sequences. G3 GenesGenomesGenetics 1:171–182.

Daguin-Thiébaut, C., S. Ruault, C. Roby, T. Broquet, F. Viard, and A. Brelsford. 2021. Construction of individual ddRAD libraries. protocols.io, doi: 10.17504/protocols.io.bv4tn8wn.

Ewels, P., M. Magnusson, S. Lundin, and M. Käller. 2016. MultiQC: summarize analysis results for multiple tools and samples in a single report. Bioinformatics 32:3047–3048.

Gutenkunst, R. N., R. D. Hernandez, S. H. Williamson, and C. D. Bustamante. 2009. Inferring the Joint Demographic History of Multiple Populations from Multidimensional SNP Frequency Data. Plos Genet. 5.

Hudson, R. R. 2002. Generating samples under a Wright-Fisher neutral model of genetic variation. Bioinformatics 18:337–338.

Martin, M. 2011. Cutadapt removes adapter sequences from high-throughput sequencing reads. EMBnet.journal 17:10–12.

Peterson, B. K., J. N. Weber, E. H. Kay, H. S. Fisher, and H. E. Hoekstra. 2012. Double Digest RADseq: An Inexpensive Method for De Novo SNP Discovery and Genotyping in Model and Non-Model Species. PLoS ONE 7:e37135.

Pickrell, J. K., and J. K. Pritchard. 2012. Inference of Population Splits and Mixtures from Genome-Wide Allele Frequency Data. PLoS Genet. 8:e1002967.

Prunus, G. 1968. Etude de systématique des populations chez l'isopode *Jaera* (*albifrons*) *albifrons* Forsman. Arch. Zool. Expérimentale Générale 109:643–702.

R Core Team. 2019. R: A language and environment for statistical computing. R Foundation for Statistical Computing., Vienna, Austria. URL http://www.R-project.org.

Rastas, P. 2017. Lep-MAP3: robust linkage mapping even for low-coverage whole genome sequencing data. Bioinformatics 33:3726–3732.

Ribardière, A., E. Pabion, J. Coudret, C. Daguin-Thiébaut, C. Houbin, S. Loisel, S. Henry, and T. Broquet. 2021. Sexual isolation with and without ecological isolation in marine isopods Jaera albifrons and J. praehirsuta. J. Evol. Biol. 34:33–48.

Rougeux, C., L. Bernatchez, and P.-A. Gagnaire. 2017. Modeling the Multiple Facets of Speciationwith-Gene-Flow toward Inferring the Divergence History of Lake Whitefish Species Pairs (Coregonus clupeaformis). Genome Biol. Evol. 9:2057–2074.

Van Ooijen, J. W. 2006. JoinMap4, Software for the calculation of genetic linkage maps in experimental populations. Kyazma B. V., Wagenigen, Netherlands.

Zhang, H. E., P. Meltzer, and S. Davis. 2013. RCircos: an R package for Circos 2D track plots. BMC Bioinformatics 14.
